# DNA metabarcoding successfully quantifies relative abundances of arthropod taxa in songbird diets: a validation study using camera-recorded diets

**DOI:** 10.1101/2020.11.26.399535

**Authors:** Yvonne I. Verkuil, Marion Nicolaus, Richard Ubels, Maurine M. Dietz, Jelmer M. Samplonius, Annabet Galema, Kim Kiekebos, Peter de Knijff, Christiaan Both

**Affiliations:** Conservation Ecology Group, Groningen Institute for Evolutionary Life Sciences (GELIFES), University of Groningen, PO Box 11103, 9700 CC Groningen, The Netherlands; Groningen Institute for Evolutionary Life Sciences (GELIFES), University of Groningen, PO Box 11103, 9700 CC Groningen, The Netherlands; Institute of Evolutionary Biology, University of Edinburgh, EH9 3FL, Edinburgh, United Kingdom; Department of Human Genetics, Leiden University Medical Centre, Leiden, The Netherlands

**Keywords:** arthropods, DNA barcoding, COI primers, insectivorous diet validation, PCR-based, Illumina sequencing

## Abstract

1. Ecological research is often hampered by the inability to quantify animal diets. Large-scale changes in arthropod diversity, abundance and phenologies urge the need to understand the consequences for trophic interactions. Diet composition of insectivorous predators can be tracked through DNA metabarcoding of faecal samples, but to validate the quantitative accuracy of metabarcoding, validation using free-living animals for which their diet can be approximated, is needed.
2. This validation study assesses the use of DNA metabarcoding in quantifying diets of an insectivorous songbird. Using camera footage, we documented food items delivered to nestling Pied Flycatchers *Ficedula hypoleuca*, and subsequently sequenced the Cytochrome Oxidase subunit I (COI) in their faeces. Our special interest was to retrieve the relative contribution of arthropod taxa with a PCR-based protocol.
3. Assessment of taxonomic coverage of the invertebrate COI primers LCO1490 and HCO1777, previously applied in insectivorous songbirds, demonstrated that COI barcodes were predominantly assigned to arthropod taxa, however, substantial amounts of reads (2–60%) were assigned to flycatchers. Modified primers reduced vertebrate reads to 0.03%, and yielded more spider DNA without significant changing the recovery of other arthropod taxa.
4. To explore digestive biases, contents of stomachs and lower intestines were compared in eight adult male flycatchers, victims of competitors for nest boxes. Similarity in arthropod community composition between stomach and intestines confirmed limited digestive bias.
5. With nest box cameras in 2013, 2015 and 2016, size-adjusted counts of prey items fed to nestlings were recorded, to approximate provided biomass of arthropod orders and families which allowed comparison with abundance of COI barcode reads in nestling faeces. The relative abundances of size-adjusted prey counts and COI reads correlated at R = 0.85 (CI:0.68-0.94) at order level and at R=0.75 (CI:0.67-0.82) at family level. Each common order and all common taxa within Lepidoptera, Diptera and Coleoptera were retrieved in similar proportions, while the taxonomic resolution was higher in barcodes than in camera records.
6. This DNA metabarcoding validation demonstrates that quantitative arthropod diet analyses is possible in songbirds. We discuss the ecological application of our approach in approximating the arthropod diets in insectivorous birds.

## 1 INTRODUCTION

Since its foundation, animal ecology has had a major focus on food (Elton, 1927): population abundances are often determined by food availability, and interactions among species are between predator and prey (or parasites and host), or predators competing for the same prey. However for generalist species with complex diets such as insectivorous birds, quantifying what an individual or a population consumes is a complicated task. Yet we need quantitative methods to characterize diets to address many ecological questions, such as: (1) how do species separate their trophic niches in space and time?; (2) what are the consequences of the global decline in arthropods for various insectivorous species within food-webs?; and (3) how do differential changes in phenology in response to global warming affect selection on breeding phenology? Influential papers on the latter two topics mostly have used correlations between features of populations, species or individual phenotypes, and general indices of food availability (e.g. Both, Bouwhuis, Lessells, & Visser, 2006; Hallmann, Foppen, Van Turnhout, De Kroon, & Jongejans, 2014), often without specifying the intermediate mechanistic link with diet. When diets are examined (e.g. Cholewa & Wesołowski, 2011; Samplonius, Kappers, Brands, & Both, 2016) this is mostly restricted to life stages when it is most easily monitored (i.e. nestlings), and, ignoring essential ecological and evolutionary features of insectivorous predators in most life and annual stages.

DNA metabarcoding (hereafter metabarcoding) can be an important tool in ecological studies (Alberdi et al., 2019), as it allows prey detection from faecal samples and thus the establishment of longitudinal and spatial studies on trophic interactions and associated biodiversity (reviewed by Valentini et al. 2009 Kress et al. 2015; and demonstrated for an Arctic foodweb by Wirta et al. 2015). Through metabarcoding, prey taxa that are visually difficult to detect can be assessed (e.g. Ando et al. 2013), new prey taxa and feeding habitats can be discovered in relatively well-studied species (Gerwing, Kim, Hamilton, Barbeau, & Addison, 2016; Trevelline et al., 2018), and ecological communities can be phylogenetically described (e.g. Evans et al. 2016). Several studies have already successfully used DNA barcodes to study insectivorous diets of bats (Zeale et al. 2011; Krüger et al. 2014; metabarcoding studies reviewed in Deagle et al. 2019) and birds (King, Symondson, & Thomas, 2015; Wong, Chiu, Sun, Hong, & Kuo, 2015; McClenaghan, Nol, & Kerr, 2019; Rytkönen et al., 2019; Shutt et al., 2020).

Relatively cheap PCR-based sequencing protocols make metabarcoding an accessible tool for ecologists, provided that PCR primers matching a sufficient reference database are available (Taberlet, Bonin, Zinger, & Coissac, 2018). For arthropods the potential of PCR-based protocols to asses diversity is emphasized by the high detection rates of 80-90% in studies using a mix of known arthropod species (Brandon-Mong et al., 2015; Elbrecht & Leese, 2015; Krehenwinkel, Kennedy, Rueda, Lam, & Gillespie, 2018; Jusino et al., 2019). In particular metabarcoding of the mitochondrial Cytochrome Oxidase subunit I (COI) is promising as it was capable of retrieving every species in the assembled arthropod communities (Krehenwinkel et al., 2018; Jusino et al., 2019), and high quality DNA barcodes of museum reference collections are available (Hebert, Ratnasingham, & DeWaard, 2003). The potential of COI has also been demonstrated using faeces of birds. Five different classes of Arthropoda and Mollusca were detected in faeces of Western Bluebird *Sialia mexicana* nestlings, using generic metazoan COI primers (Folmer, Black, Hoeh, Lutz, & Vrijenhoek, 1994) that amplify 710 bp of the COI gene (Jedlicka, Sharma, & Almeida, 2013), but this long fragment may not recover all arthropod taxa (Jusino et al., 2019). Additionally, in several warblers and Barn Swallows *Hirundo rustica* it was shown that using a shorter COI fragment may result in more PCR product and more arthropod taxa being recovered from avian faeces (King et al., 2015; McClenaghan et al., 2019; Rytkönen et al., 2019). These are encouraging results for ecologists interested in metabarcoding arthropod diets from faecal samples. However, a number of issues persist.

Most importantly, it is unknown if metabarcoding can result in a quantitative assessment of arthropod diets (Deagle et al., 2019). Many studies assess the presence/absence of taxa and not the read abundance, because with PCR-based methods may not sufficiently approximate the relative abundance of each prey taxa (Elbrecht and Leese 2015; Piñol et al. 2015; Jusino et al. 2019, but see Thomas et al. 2016; Deagle et al. 2019). Therefore, validation tests of birds fed with recorded food items are important, especially for generalist species with more diverse diets (King, Read, Traugott, & Symondson, 2008; Pompanon et al., 2012). Also, potential biases due to differences in how taxa pass through the digestive track need to be assessed (King et al., 2008). And more technically, studies have reported PCR inhibition due to uric acids in avian faeces which leads to loss of samples and jeopardizes study design (Jedlicka et al., 2013; Rytkönen et al., 2019), and difficulties in retrieving all arthropod taxa with a single PCR protocol, especially due to PCR primers mismatches with spiders (Jusino et al., 2019).

This paper aims to examine whether metabarcoding can provide a quantitative estimate of the relative contribution of taxa to the diet, or whether it is restricted to a qualitative assessment of the frequency of occurrence of prey in samples, while addressing the technical pitfalls. We provide methods on how to (1) optimize DNA-extraction from avian faeces and (2) maximize reads of arthropod taxa with adjusted primers. We show (3) that there is little differential loss of diet items throughout the digestive track, and validate (4) that there is a good quantitative match between approximated diets and the relative read number of diet taxa. Our study population of Pied Flycatchers *Ficedula hypoleuca* gives the opportunity for a validation study in a natural setting. Adult birds breed in nest boxes and provide their nestling with a large variety of taxa which is recorded on camera (Samplonius et al., 2016; Nicolaus, Barrault, & Both, 2019) and nestling faeces can be easily collected.

## 2 MATERIAL AND METHODS

### 2.1 Study design

We tested the application of massive parallel sequencing, or metabarcoding, of the mitochondrial gene COI to quantify arthropod diets in birds by means of three methodological steps followed by a final validation test.

1. In Step 1, we explored the full extent of taxa that could be detected in DNA templates extracted from faecal samples, by deep-sequencing PCR products obtained with published generic COI primers targeting arthropods (King et al., 2015). For this, faecal samples of Pied Flycatchers adults (n = 3) and nestlings (n = 2) were sequenced to a depth of >2 million reads per sample.
2. In Step 2, we evaluated (a) DNA-extraction methods, (b) primer pairs to exclude host DNA while avoiding blocking primers (Piñol, San Andrés, Clare, Mir, & Symondson, 2014) and (c) sequencing depth. In a double pair-wise design, faecal samples of two nestlings were divided over two DNA-extraction methods, and for each extraction method two different PCR primer pairs were tested, allowing pair-wise comparisons of methods. The samples were sequenced in two runs: a limited run aiming for 10,000 reads per sample and an extended run aiming for >50,000 reads per sample.
3. In Step 3, we explored possible effects of digestion on prey taxa detection and prey community, by metabarcoding the gizzard and lower intestines content of adult Pied Flycatchers (n = 8).
4. In the final validation test, we compared arthropod taxa detected in faeces of Pied Flycatcher nestlings (n = 63) with the prey taxa provided by their parents, as determined through camera observations of prey items delivered to the nest box (n = 39 observation days in three years). Analysing this test, we also tested pipelines settings.

### 2.2 Field faeces collection and camera observations on diet

The five faecal samples for Step 1 were collected in spring 2013 in the study area in Drenthe, The Netherlands (n =3; 52°49’ N, 6°25’E; see Both, Bijlsma, & Ouwehand (2016) for detailed description), and in 2011 in winter in Ghana (n = 2; 7°58’N, 1°44’ W; J. Ouwehand, pers. comm.). Samples for Step 2–4 were collected in Drenthe. For Step 2, faeces were collected in May 2015 from two chicks of two nests and at two ages, namely 3 and 7 days old. For Step 3, we used eight male adult Pied Flycatchers that were killed by Great Tits *Parus major* in a nest box between 14 April and 17 May 2015, a consequence of heterospecific competition for nest boxes (Slagsvold, 1975; Merilä & Wiggins, 1995; Samplonius & Both, 2019). For the validation test, faeces were collected from chicks whose food provisioning by their parents was monitored by cameras fitted inside the nest boxes. In 2013 and 2015, faecal samples of 1–3 chicks per nest box were collected on the same day or one day before or after the camera recording day. We allowed this range of days because provisioning behaviour is repeatable between subsequent days (Nicolaus et al., 2019). However, to test if more targeted timing is important for validation, in 2016 faeces of two chicks per nest box were always collected on the same day at the end of the recording period. Samples were placed in a sterile 2.0 mL tube with 96% ETOH and stored at −20 °C, except in Step 2 samples were flash-frozen and stored at −80 °C in PBS buffer. For long-term storage all samples were stored in −80 °C freezers.

From 39 camera sessions of ca. 2 hours (five in 2013, 18 in 2015, and 16 in 2016) camera footage was scored, noting the food item provided and its relative size in relationship to the beak of the adult (details in Samplonius et al. 2016). To allow comparison with the read counts obtained from metabarcoding of faeces which reflect ingested biomass and not prey numbers, the prey counts from camera footage were size-adjusted to approximate biomass. With this adjustment large prey items were given extra weight compared to small prey, by using a multiplication factor varying from 0.04 to 6.0 depending on the prey size relative to the bill size. Prey counts were taxonomically assigned to arthropod class, order, and if possible also to family, genus and species, yielding taxonomically unique groups (for comparison with COI data called “camera-OTU”) that could vary in precision of taxonomic assignment. The camera-OTU “unknown” accumulated all counts of unclassified animals. Prey items were taxonomically assigned without prior knowledge of metabarcoding results, and no prey items were reassigned *a posteriori*.

### 2.3 DNA-extractions

In Step 1, complete faecal samples were used. In Step 2, each faecal sample was homogenized and split in two subsamples for each extraction method. In Step 3 and 4, samples were subsampled to arrive at a sample weight of < 1 g to reduce levels of uric acids, which are present in bird faeces and cause PCR inhibition (Jedlicka et al., 2013). Subsamples were taken using a small metal scoop sterilized at the flame. Before proceeding with the extraction, samples were placed at 55 °C for 10-20 min until all ETOH was evaporated. DNA was extracted with Qiagen DNeasy PowerSoil Kit, formely made by MoBio (Step 1), a kit recommended for bird faeces (Vo & Jedlicka, 2014), and with the Invitrogen™ PureLink™ Microbiome DNA Purification Kit (Step 3 and 4). The PureLink kit would result in less uric acid and higher PCR success. In Step 2 the two kits were compared.

To increase the yield of prey DNA, the manufacturer’s protocols of both kits were altered as follow: (1) a bead beater was used instead of a vortex mixer, in 3×1 min bouts (Powersoil), or 5×2 min bouts (PureLink) pausing 30 sec between bouts, (2) approx. 0.1 g extra 0.1 mm Zircona/Silica beads were added, (3) for the final elution we used 20 μl Ambion© purified DNA-free water, and (4) pre-elution incubation was extended to 4-5 min and DNA was re-applied to the filter and incubated 2 min extra before final elution.

To prevent contamination, arthropod specimen were not allowed in our lab. As a consequence metabarcoding of reference arthropod material from Africa (not reported here) was performed at another institute. All materials used were autoclaved and UV-sterilized for 20 min. Clean gloves were used for each sample. Two negative control extractions with no faecal sample were included to test the purity of the extraction kits as well as make sure there was no contamination between samples during the extraction procedure. DNA concentrations were not normalized before PCR because faecal samples contain more bacterial and host DNA than target prey DNA.

### 2.4 PCR primers

Primer choice was evaluated (Alberdi, Aizpurua, Gilbert, & Bohmann, 2018) as follows. In Step 1, following the successful assessment application in insectivorous songbirds (King et al., 2015) we used the generic invertebrate COI primers LCO1490 (Folmer et al., 1994) and HCO1777 (Brown, Jarman, & Symondson, 2012) (Table S2-1). For Step 2, the forward primer and two altered reverse primers were created, introducing miss-priming with flycatcher DNA but improving the match with arthropod DNA. To design the altered primers an alignment of host DNA and various prey taxa was constructed in Geneious 9.0.5 using GenBank sequences of *Ficedula sp.*, *Araneae, Coleoptera, Diptera*, *Hemiptera*, *Hymenoptera* and *Lepidoptera* (Table S2-2). We tested which combination of primers successfully amplified faecal DNA but avoided host DNA, using flycatcher DNA (obtained from a molecular sexing project, M. van der Velde, pers. comm.) as positive control. In Step 3 and 4 the modified primers LCO1490_5T (5’-GGTCTACAAATCATAAAGATATTGG-3’) and HCO1777_15T (5’-ACTTATATTATTTATACGAGGGAA-3’) were used.

### 2.5 PCR conditions

PCR reactions were set up in a DNA-free room. Each sample was done in duplicate or triplicate to avoid PCR bias, and from each PCR master mix negative controls were taken to track possible contamination of PCR reagents. Before pooling, negative controls were assessed with 5 μl PCR product in a standard gel electrophoreses. Annealing temperature was set low to minimize taxonomic bias (following (Ishii & Fukui, 2001)).

PCR reactions had a final reaction volume of 20 μl containing 2.5 μl 10x Roche buffer, 0.2 μl 25 mM dNTPs, 0.88 μl 50 mM MgCl2, 0.03 μl BSA, 1.0 μl of each primer (10 μM), 0.2 μl 5U/μL Taq polymerase (Roche) and 5 μl DNA template. The PCR profile included an initial denaturation at 94°C for 2.5 min., 35 cycles of 94°C for 30 s, 48°C for 30 s and 72 °C for 45 s, and a final extension at 72°C for 10 min. In Step 3, for eight of 16 samples AccuStart II PCR ToughMix© was used to improve the amplification success. In this test DNA may have been degraded, as samples were taken from birds that may have been dead for a day before they were found in the nest box. The reaction volume was 10 μl including 5 μl AccuStart, 1μl of each primer (10 μM), 1μl ddH2O and 2μl DNA template. When using AccuStart the PCR profile was altered to 3 min. at 94 °C followed by 35 cycles of 1 min at 94 °C, 30 sec. at 48 °C and 1 min. at 72 °C, and a final extension at 72°C for 10 min.

### 2.6 Massive parallel sequencing

PCR products of each test (total *N* = 5+8+18+63 = 94) and the pooled negative extraction controls (*N* = 2) were sequenced on the MiSeq© Sequencer (Illumina) at the Department of Human Genetics, Leiden University Medical Centre. Libraries were prepared with the MiSeq© V3 kit, generating 300-bp paired-end reads. The V3-kit does not normalize, which means that it leaves the relative presence of initial PCR product intact, and therefore this library preparation method allows assessing the relative contribution of prey taxa.

### 2.7 Pipeline design: empirical selection of settings

Firstly, using the software USearch 9.2 (Edgar, 2010) for each test separately we extracted unique high-quality barcode reads (molecular operational taxonomic units, abbreviated as OTU) with seven command lines. If applicable, settings were empirically tested (Alberdi et al., 2018).

(1) Paired reads (the forward and reverse reads of 299 bp each) were merged, and to obtain a consensus sequence of 281 bp. This removed the unaligned segments at both ends which contain the sequencing adaptors.
(2) Primer sequences were removed by truncating each end by 25 bp, the length of the longest PCR primer. The 25 bp was determined by visual inspection of the merged paired reads in Geneious indicating the previous algorithm had not sufficiently removed primer sequences.
(3) Reads were filtered for quality at a default error (E) value of 0.4, where E = 1 means all reads incl. low quality reads are included, and decreasing E values means more stringent filtering. Reads were truncated to 220 bp with the same command line. This size was determined by visual inspection of reads after running command line 2. For the validation test, these two settings were evaluated (see *Data analyses*).
(5) Next, reads were de-replicated by finding duplicated reads and assigning a count to unique reads. This was necessary to merge identical reads that are present in both orientations. Subsequently, the singletons were removed.
(6) Using the UPARSE-OTU algorithm (Edgar, 2010) reads that were minimally 97% identical were clustered and the consensus sequence of each cluster was assigned an OTU ID; this created an OTU sequence database. This algorithm also filters chimeras.
(7) Lastly, for each sample the number of reads (paired and with primers truncated) that matched with each OTU was determined, resulting in an OTU frequency table. The default identity match of 97% was used. This setting was evaluated by comparing OTU tables created with 90% and 97% identity matches in the validation test; as values highly correlated, we believe that more stringent matching with a 97% cut-off was possible without data loss.

In Step 3 and 4, the final OTU frequency table was adjusted for the pooled negative extraction and PCR controls, by deducting the number of reads found for an OTU in the pooled negative extraction controls from each cell in the OTU table. The sum of reads in the pooled negative controls was 70 reads with a maximum of 10 per OTU (Table S3-1).

### 2.8 Taxonomic assignments

The obtained OTU databases were searched against the *nr* database in GenBank (Benson, Karsch-mizrachi, Lipman, Ostell, & Sayers, 2009), using the BLAST function in Geneious 8.1.7 (Kearse et al., 2012). We used GenBank, which also contains the public part of sequences from BOLD (Barcode of Life Data Systems) (Ratnasingham & Hebert, 2007), because species diversity of the Western European arthropods is sufficiently covered (King et al., 2008). Also, validating our approach with a public database will demonstrate the general applicability for other European studies.

We used the Megablast option which is faster than blast-n and only finds matches with high similarity. Settings were: max e-value = 1e-1 (the lower the number expected (e) hits of similar quality the more likely the hit is real), word size = 28 bp (minimal match region) and gap cost = linear. The best hits were saved in a query-centric alignment. For each match between an OTU and a reference organism we recorded: non-annotated matching sequences, query coverage, bit-score, e-value, pairwise identity, sequence length and grade. The grade is a percentage calculated combining three statistics: the query coverage, e-value and pairwise identity values for each hit, which have a weight in the equation of 0.5, 0.25 and 0.25, respectively. Each encountered reference organism was included in a taxonomy database with its GenBank Accession number. Inclusion of a reference organism in our taxonomy database was independent of the likelihood of occurrence in the study area (i.e. in some cases an OTU matched best with a species not occurring in Europe).

Taxonomic categories included were Kingdom, Phylum, Class, Order, Family, Genus and Species. The utility of assignment to the order and family level were explored in the validation test. Digestive biases (Step 3) were assessed at the genus level. The grade score was used to assess the reliability of taxonomic assignment. We considered species assignments only indicative of the actual species. In general, species assignments through a similarity match of short OTU reads to reference sequences is ambiguous: even when an OTU has a 100% match with a species barcode, the probability that it is the same species is not 100% (Ward, 2009).

### 2.9 Data analyses

Data analyses were performed in *R*, using packages *phyloseq* (McMurdie & Holmes, 2013), *car* (Fox, Friendly, & Weisberg, 2013) and *vegan* (Dixon, 2003). Read counts of taxa in each sample were transformed to relative read abundance (RRA) on the order or family level. To arrive at a higher aggregate scale (e.g. year), average RRA (±SD) per taxa per sample were calculated (step 1: percentage per sample, step 2: average across these percentages). Frequency of occurrence of taxa (FOO) was expressed as the number of samples in which taxa occurred. Arthropod community differences were assessed with ordination analyses, applying nonmetric multidimensional scaling (NMDS) using the Bray-Curtis distance (Anderson, 2001).

In Step 1 and 2 the contribution of all kingdoms and phyla was assessed to explore amplification of non-target taxa and possible contaminants. Subsequently in Step 2-4, the data was pruned to the target phylum Arthropoda. The effect of read depth (sample quality) on the detected variation of arthropod OTUs was assessed with Pearson’s correlation tests using various diversity indices. In the camera records, to create diversity plots the size-adjusted counts had to be rounded to integers.

In Step 2, the linear fit between retrieved abundances of arthropod orders across laboratory methods was explored by least square regressions.

In Step 3, gizzard-intestine differences in the FOO of genera were tested with a contingency Chi square analyses; this analyses was restricted to common taxa, occurring in =>3 samples. NMDS ordination was performed in *phyloseq* on genus level (because at an average grade of 99% taxonomic assignments were sufficiently reliable) with categorical variables sample type and bird ID; read counts per sample per genera were normalized to median count, and singleton genera were pruned to reach convergence. To assess the degree of difference between groups a *permutest* was conducted.

In the validation test, to evaluate which metric better described the camera-recorded diet, FOO or RRA, Pearson’s correlation tests were applied. In contrast to rank correlations, this test allows to explore differences in the linearity of relationships quantitatively. The variation in the correlations between single faeces/camera-session combinations was not explored with multilevel models because (1) to include FOO in the comparison aggregated data had to be used, (2) no predictor values exists because of uncertainties in taxa abundance estimates from camera footage. For this analyses the parasites orders Mesostigmata, Prostigmata, Sarcoptiformes, Siphonaptera, Trombidiformes and the Braconidae, Eulophidae, Ichneumonidae families in Hymenoptera were excluded. NMDS was performed on order and family level in *vegan* with wrapper function *metaMDS*, which allows using prey taxa proportions (Oksanen et al., 2019), followed by *permutest*. Categorical variables were sample type, year, camera session ID; date was included as numerical variable. To assess the effect of rare taxa, ordination of full data sets was compared to pruned datasets. Orders and family were pruned an RRA =>1% (calculated over all samples). For families, proportions were max-transformed with command *decostand*, meaning all proportions were standardized to the family with the highest sum of proportions.

Also in the validation test, pipeline variants were tested on pooled data of five samples (T0113, T0218, T0313, T0420 and T0520) to assess effects of quality filtering and read truncating. We tested 18 settings: E-values of 0.1–1.0 (while truncating at 220 bp), and truncation values of 140–280 bp (while filtering at 0.4). We expected that (a) stringent quality filtering reduces the number of detected species, and (b) intermediate trimming settings yields most species, because OTUs based on short reads have poorer BLAST results while OTUs based on long reads results in data loss. BLAST searches were run independently for each pipeline variant, and OTUs were collapsed to species, meaning that reads of OTU variants assigned to the same species were summed. The pipeline variants were assessed for all taxa, and for a 99% dataset created by cumulatively removing rare species until 99% of the reads were left (this excluded species with <0.01% reads). Differences between pipelines were statistically tested by a One-Way Repeated Measures ANOVA on number of reads (abundance) per arthropod family, using the single Anova(mod, idata, idesign) function in *car*. The dependent variable *mod* was the linear model correlating reads per family between the 18 variants, *idate* was our dataframe Family abundance, and the factor *idesign* was our pipeline variant.

## 3 RESULTS

### 3.1 Step 1: retrieval of target phylum Arthropoda

The five test samples yielded 922 OTUs representing 5.3 million reads of which 368 OTUs (representing 3.3 million reads) were assigned to the target phylum Arthropoda. Arthropod OTUs were divided over five classes, 25 orders and 84 families and 122 genera (Table 1).

**TABLE 1.** Overview of the data obtained in the four stages of the study, including samples sizes, PCR reactions, read abundances and number of OTUs. In Step 3, the number of reactions was 17 because a PCR replicate for one gizzard sample was included. In the validation test, both DNA barcodes and camera observations were used; for the latter size-adjusted prey counts are listed instead of barcode reads. (*) In the validation test, COI data contained 5 arthropod classes of which 4 matched the classes detected in the camera records. (**) In Table 2 the number of orders is 23 because singletons were excluded. V = validation test. NC = negative control.

| Test | Data type | Sample size: birds/nests (PCR reactions) | Age birds | Total assigned reads | N OTUs | Reads after subtraction NC | OTUs after subtraction of NC | Reads in Arthropoda | Arthropod OTUs | Median reads in samples | N classes* | N orders** | N families | N genera |
| --- | --- | --- | --- | --- | --- | --- | --- | --- | --- | --- | --- | --- | --- | --- |
| 1 | COI barcode | 5 (5) | all | 5,328,506 | 922 | - | - | 3,275,344 | 368 | 460,528 | 5 | 25 (23) | 84 | 122 |
| 2 | COI barcode | 2 (8) | nestlings | 219,241 | 424 | - | - | 192,228 | 244 | 25,112 | 5 | 17 | 64 | 101 |
| 3 | COI barcode | 8 (17) | adults | 887,188 | 260 | 886,749 | 258 | 797,422 | 180 | 46,127 | 3 | 12 | 53 | 81 |
| V | COI barcode | 63 (63) | nestlings | 912,130 | 1,083 | 911,947 | 1,018 | 897,315 | 832 | 9,119 | 5(4) | 24 | 145 | 297 |
| V | Camera | 39 (NA) | nestlings | - | - | - | - | 10,458 | 124 | 176 | 4 | 18 | 59 | 40 |

In the African samples 64-159 OTUs and 11 arthropod orders were detected, while the Dutch samples yielded 326-412 OTUs and 17-21 arthropod orders (Table 2). OTUs that could not be taxonomically assigned represented 134,866 reads (2.5%; range 0.27%–5.58%) (Fig. S1-1A).

**TABLE 2.** Overview of the yield of reads, OTUs and assignment success in Step 1, using published arthropod primers LCO1490 (Folmer et al., 1994) and HCO1777 (Brown et al., 2012). The test included five samples from The Netherlands and Ghana - West Africa. For each barcode ID the life stage of the associated sample is given.

| BarcodeID - life stage | N raw reads | N reads to OTUs | N reads to Aves | % reads to Aves | N OTUs | Arthropoda orders | PCR product | Origin |
| --- | --- | --- | --- | --- | --- | --- | --- | --- |
| T1118 - Adult | 3,107,761 | 1,726,331 | 136,794 | 8% | 339 | 17 | high quality | Netherlands |
| T1318 - Chick | 2,716,379 | 1,338,868 | 257,454 | 19% | 326 | 17 | high quality | Netherlands |
| T1218 - Chick | 2,219,834 | 836,877 | 496,821 | 59% | 412 | 21 | midrange quality | Netherlands |
| T1018 - Adult | 2,608,173 | 1,227,054 | 27,896 | 2% | 159 | 11 | high quality | Africa |
| T0918 - Adult | 2,014,607 | 199,376 | 7,583 | 4% | 64 | 11 | low quality | Africa |
| **TOTAL** | **12,666,754** | **5,328,506** | **926,548** | **17%** | **922** | **23** |  |  |

Arthropod OTUs were mostly Insecta (2.8 million reads) and Arachnida (0.5 million reads) (Fig. S1-1C). In the sample that yielded low quality PCR product, as assessed through gel electrophoreses, many raw reads were not assigned to OTUs, but this had no effect on the number of detected arthropod orders (Table 2).

OTUs assigned to non-target groups such as plants and fungi consumed 1.1 million reads, but not all samples had significant numbers of non-target reads (Fig. S1-1A). Of the 4.2 million reads assigned to Animalia, 932,551 reads were Chordata, mostly flycatcher DNA (926,548 reads) (Fig. S1-1B, D). Host DNA reads were especially abundant in nestlings whose faeces are wrapped in a faecal sac and took up 2.3–59.4% of the reads (Table 2).

**In summary, this initial step revealed the need to adjust protocols to increase the yield of target taxa and avoid amplification of host DNA.**

### 3.2 Step 2: primer redesign, DNA-extraction method and read depth

In Step 2, 219,241 paired reads were obtained assigned to 424 OTUs (Table 1). Animalia had 205,603 reads of which 192,228 were Arthropoda, assigned to 244 OTUs, divided over 5 classes, 17 orders and 64 families and 101 genera. Chordata (Aves, Mammalia) had 6.5% of the reads (details in Fig. S2-1). The median number of arthropod reads per PCR product was 25,112 (range 15,133-36,564; merged runs).

In the four PCR replicates, the RRA of arthropod orders correlated between the original and modified primers at R^2^ = 0.98, R^2^ = 0.98, R^2^ = 0.72 and R^2^ = 0.99, respectively (Fig. 1; Table S2-3). The number of reads “lost” to Chordata (mostly Aves) decreased from overall 12,933 reads (6%) with the original primers, to 3 reads (0.001%) with the modified primers (Fig. S2-1B). The number of reads of mammalian DNA decreased from 0.15% to 0.03% (Fig. S2-1C). Although overall no differences were observed between primers in the RRA of arthropod orders (Fig. 1; R^2^ = 0.92, p < 0.001), the modified primers yielded more Araneae reads (5.6%) than the original primers (1.7%). Also in Hemiptera (true bugs) and Hymenoptera (mostly ants and some parasitoid wasps) there were small differences in the same direction between all four replicates (Table S2-3).

**FIGURE 1.**
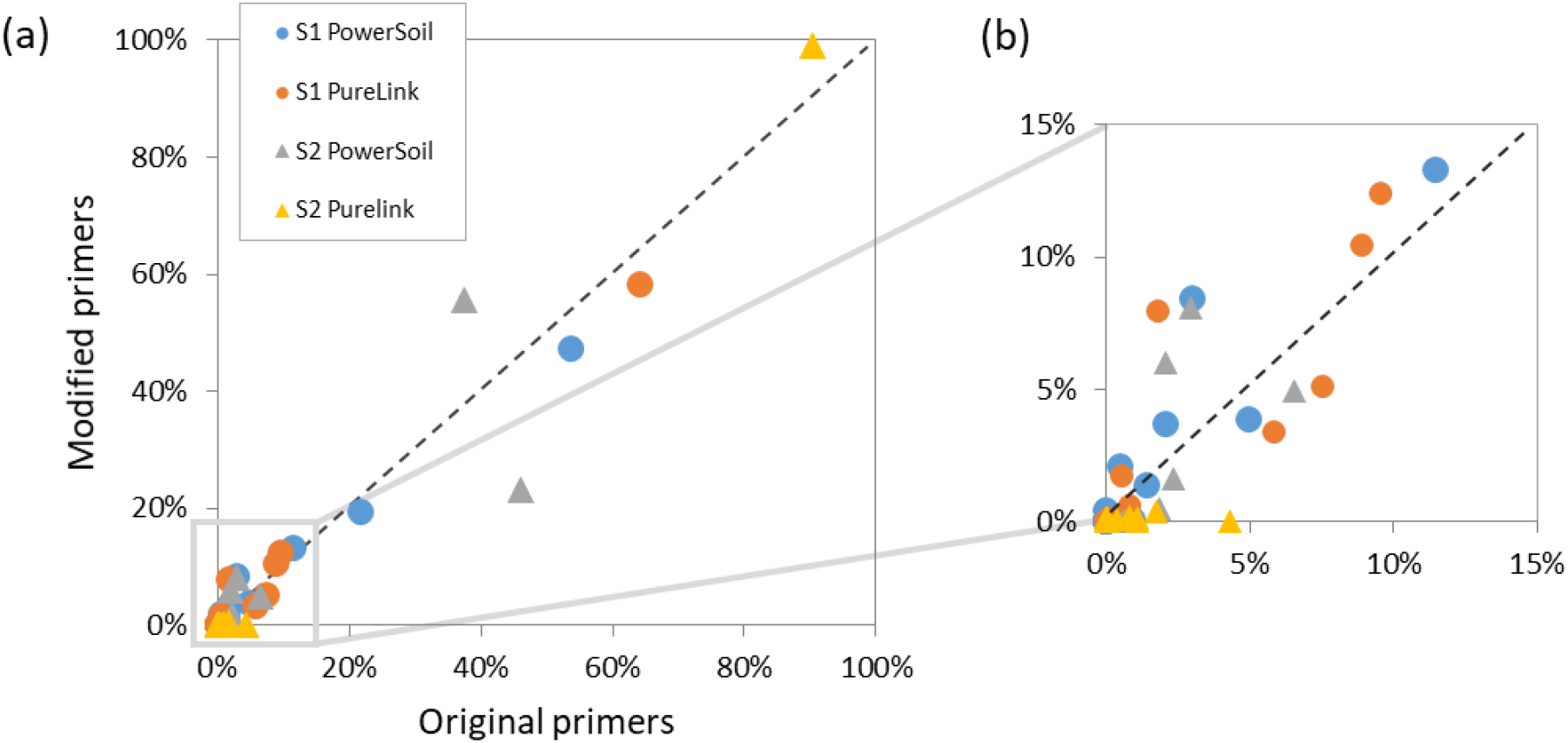
Pairwise comparison of original versus modified primers tested in Step 2. Given are RRA (%) in each sample of each of 17 arthropod orders listed in Table S2-3. Panels shows (a) all detected orders and (b) orders represented by less than 15% reads. Dashed lines depicts X = Y. S1 and S2 are the sample IDs each subjected to two extraction methods: PowerSoil and PureLink; each DNA extraction was tested with the original and modified primers (for details see Table S2-3 and Fig. S2-1A).

DNA-extractions with PureLink yielded more total reads than PowerSoil extractions (between 5,121– 12,443 extra in four replicas, Fig. S2-1A), and contained a similar to higher proportion of reads assigned to any taxa (99.95% versus 98.77%) and to Animalia (96.3% versus 90.3%). For sample 1, the RRA of arthropod orders did not vary between extraction methods (average R^2^ = 0.94; left four columns in Table S2-3); in sample 2 the PureLink extraction was dominated by Hymenoptera (average R^2^ = 0.57; right four columns in Table S2-3). This was consistent between PCR replicates.

The sequencing depth of the replicas that were sequenced in the limited versus extended Illumina run (n = 16) was 9,777–11,908 versus 50,056–80,113 raw reads (2,400-6,700 and 16,500-36,300 paired reads). In this range no effect of read depth was found on the number of arthropod OTUs (Fig. S2-2).

**In summary, we (i) increased the yield of target COI reads by applying an alternative DNA-extraction method, (ii) reduced of amplification of non-target avian/mammalian DNA through adjustments to the PCR primers, and (iii) established target sequencing depths of 2,000-10,000 reads per sample.**

### 3.3 Step 3: digestive bias

For the eight paired gizzard and intestines samples (n = 17 including a PCR replicate), we obtained 887,188 paired reads assigned to 260 OTUs. After subtraction of the maximum number of reads found for negative control OTUs, 886,749 reads (12,171 to 92,334 reads per sample) were assigned to 258 OTUs. The target phylum Arthropoda had 797,422 reads; the second largest group were parasitic worms found in one intestine sample (phylum: Acanthocephala, Fig. S3-1A).The median number of reads per sample assigned to arthropods was 46,127 (range 11,431–91,418), divided over 12 orders, 53 families and 81 genera (Table 1). OTU richness and Shannon-diversity did not correlate with the number of reads (resp. R = 0.25 (CI −0.26-0.65); p = 0.33 and R = --0.32 (CI −0.69-0.19); p = 0.21, Fig. S3-1B). All genera had high taxonomic assignment grades of on average 99% (90-100%) with an outlier of 49% for one genus (Diploplectron).

Overall arthropod communities were more similar within an individual’s gizzard and intestines than between individuals (Fig. 2; permutest (Sample Type): F = 0.053, p = 0.82; permutest (Bird ID): F = 1.039: 21 pairwise p values ≤ 0.01, 7 pairwise p values > 0.1), although patterns varied between birds (Fig. S3-2). Also the FOO of common taxa was not significantly different between organs (X^2^ = 3.721, df = 16, p = 1.00). In both gizzards and intestines, FOO was highest for *Kleidocerys* (Hemiptera), *Boletina* (Diptera), *Strophosoma* (Coleoptera) and *Formica* (Hymenoptera) (Table S3-2). The Diptera genera *Aedes* and *Pollenia* had a slightly lower occurrence in intestines than in gizzards. PCR replicates of the gizzard of flycatcher AV82435 had very similar arthropod communities (Fig. 2).

**FIGURE 2.**
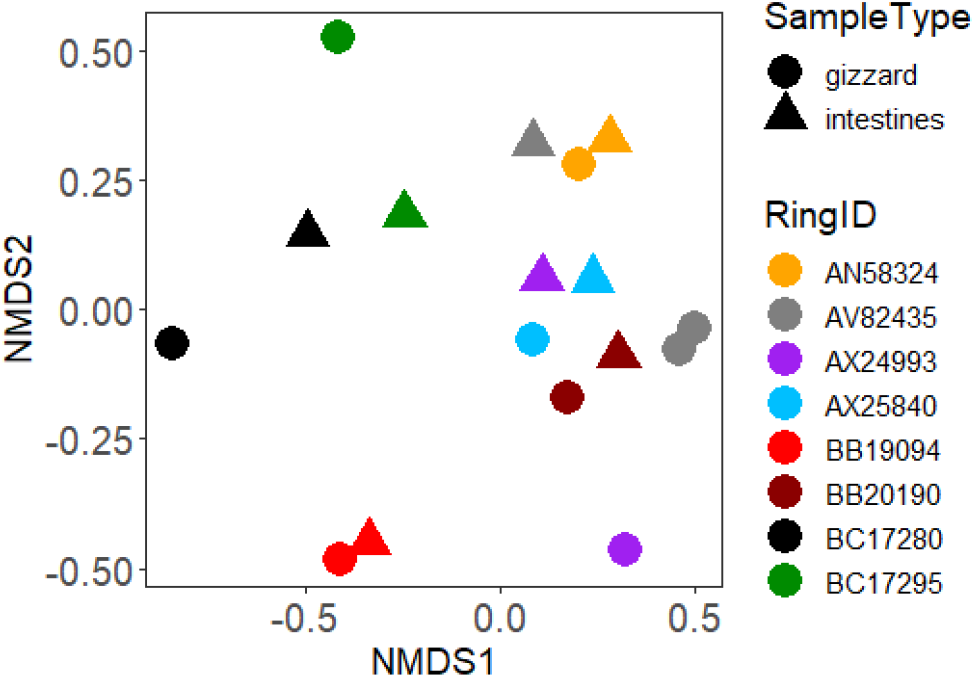
Arthropod taxa community in gizzard and intestinal samples of adult Pied Flycatchers represented by a Bray-Curtis ordination plot of paired samples types, colour-coded by each bird’s ring ID (metal band number); note that for bird AV82435 a gizzard PCR replicate is shown.

**In summary, we established little digestive bias and no loss of common taxa between gizzard and intestines.**

### 3.4 Validation test: retrieving the relative contribution of taxa

In 39 camera sessions a total of 7,314 food items were counted. Scaling in size relative to bill size had a slight effect on the relative abundance of orders, especially increasing the relative importance of Lepidoptera (Fig. S4-1). The median number of food items (”reads”) observed per camera session was 118 (range 61–468). These prey counts were divided over 124 taxonomically unique groups (Fig. S4-2A, for comparison with COI data called camera-OTU) belonging to four arthropod classes: Arachnida, Insecta, Diplopoda and Malacostraca. In total 123 camera-OTUs were identified on order level (18 orders), 105 OTUs on family level (59 families), and 46 OTUs on genus level (40 genera). One camera-OTU contained the accumulated 1,040 counts (9.9%) of “unclassified animals”. Camera-OTU richness varied per camera session (Fig. S4-3A), where observed OTUs varied with the number of reads i.e. prey counts (R = 0.36 (CI 0.05-0.61); p = 0.02; Fig. S4-3B) but Shannon-diversity did not (R = 0.31 (CI −0.01-0.57); p = 0.06; Fig. S4-3C).

A total of 4,585,185 raw reads were obtained for 63 faecal samples of nestlings (45,224–146,704 per sample). Between pipeline settings taxonomic assignment to class, order and family did not vary significantly (F_1,17_ = 0.17, p = 0.99) (Fig. S4-6A,B). The diversity indices showed a slight optimum when trimming was set to 180-220 (Fig. S4-6C,D). For further analyses therefore filtering was set at E-values of 0.4 and trimming at 220 bp. We obtained a total of 912,130 assigned reads, which after subtraction of negative control reads was reduced to 911,947. The total number of OTUs (after subtracting singletons) was 1,018.

This COI barcode data set was reduced to the four arthropod classes found in the camera records (Arachnida, Insecta, Diplopoda and Malacostraca) plus Chilopoda, representing 98.4% of the assigned reads (897,315 of 911,947) and 832 OTUs, covering 24 Arthropod orders. Overall 145 arthropod families were detected and 297 genera which had a median assignment grade of 99.9 (mean 98.2). For nine genera the grade score was below 90; they were represented by 232 reads (2-119 reads per sample). 43 genera had grades of 90-97 suggesting that the actual genus may not have been available on GenBank; six of these uncertain genera were abundant (>5,000 reads): *Panolis*, *Kleidocerys*, *Phanomorpha*, *Philoscia*, *Eridolius* and *Phyllopertha (*note that *Phanomorpha* is restricted to Australia). The three most abundant OTUs present with >50,000 reads (max. 93,244) were assigned to *Panolis flammea* (Lepidoptera, Pine beauty), *Kleidocerys resedae* (Hemiptera, birch catkin bug) and *Porcellio scabe*r (Isopoda, common rough woodlouse). The median number of all reads per sample was 9,119 (range 108 –52,573); in six samples <1000 reads were assigned (Fig. S4-2B). The read depth per sample correlated with the observed number of arthropod OTUs (R = 0.56 (CI 0.35-0.71); p <0.001), but not with Shannon-diversity (R = −0.03 (CI −0.28-0.22); p = 0.83; Fig. S4-4). In eight faecal samples >90% reads were of a single order (Fig. S4-5), mostly represented by a single genus; samples were dominated by Diptera (n = 3), Hymenoptera (n = 2), Hemiptera (n = 1), Coleoptera (n = 1) or Lepidoptera (n = 1) (Table S4-1).

To compare COI barcodes with camera records from the same broods, 4 of the 63 samples were discarded for low read quality or missing camera records, leaving a dataset of 59 faecal samples and 39 camera sessions (Table 3). For the camera sessions which had duplicate (n =18) or triplicate (n = 1) faecal samples, the variation in arthropod communities detected in the COI barcodes was larger between camera sessions than between repeated samples (permutest (Camera ID) - Order: F = 34.67, p < 0.001, Mean Sq(groups) = 0.030, Mean Sq(residuals) = 0.0009; permutest (Camera ID) - Family: F = 7.86, p < 0.001, Mean Sq(groups) = 0.007, Mean Sq(residuals) = 0.001) (Fig. 3).

**FIGURE 3.**
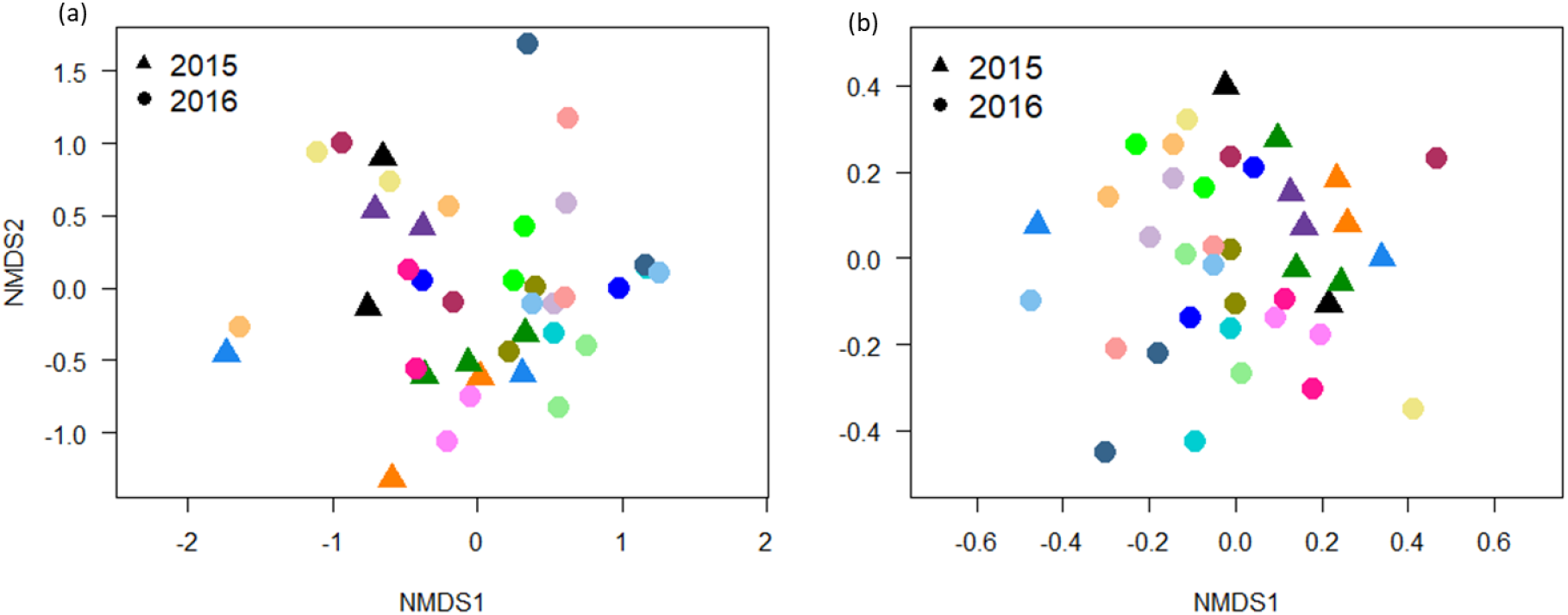
Similarity of arthropod taxa communities between COI metabarcoded faecal samples collected in the same camera session (*N* = 19). Shown are paired samples from 18 camera sessions, and three samples collected in one session (dark green triangle). Given are non-metric Bray-Curtis ordination plots for all observed orders (a) and the 50 most abundant families (b). Each cameras session is represented by a different colour, each year by a different symbol, and each COI sample by a data point.

**TABLE 3.**
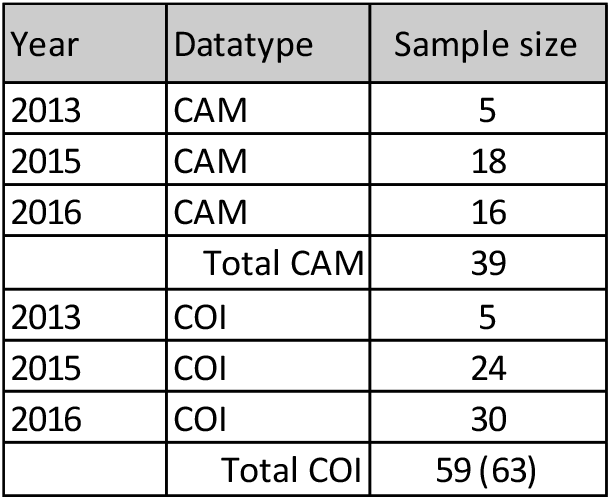
Overview of samples sizes in the validation test. Note that sample size of faeces for with COI metabarcoding data was obtained was reduced from 63 to 59 in further analyses.

The arthropod communities detected in the COI barcodes was significantly more diverse than on camera records (Fig. 4), both at the order level (Fig. S4-7a,b; F = 72.53, p < 0.001) and family level (Fig. S4-7c,d; F = 84.33, p < 0.001); note that on family level, per camera session on average 39% of the items were not assigned to family, and hence unknown. COI barcodes contained 22 orders versus 18 in the camera records. In the six most abundant orders, two times more families (105 versus 50) were detected in COI than on camera: Diptera (33 families: 31 versus 13), Lepidoptera (22: 18/9), Coleoptera (16: 15/10), Hymenoptera (15: 13/6), Hemiptera (11: 10/4), and Araneae (18: 18/8) (see also Fig. S4-8).

**FIGURE 4.**
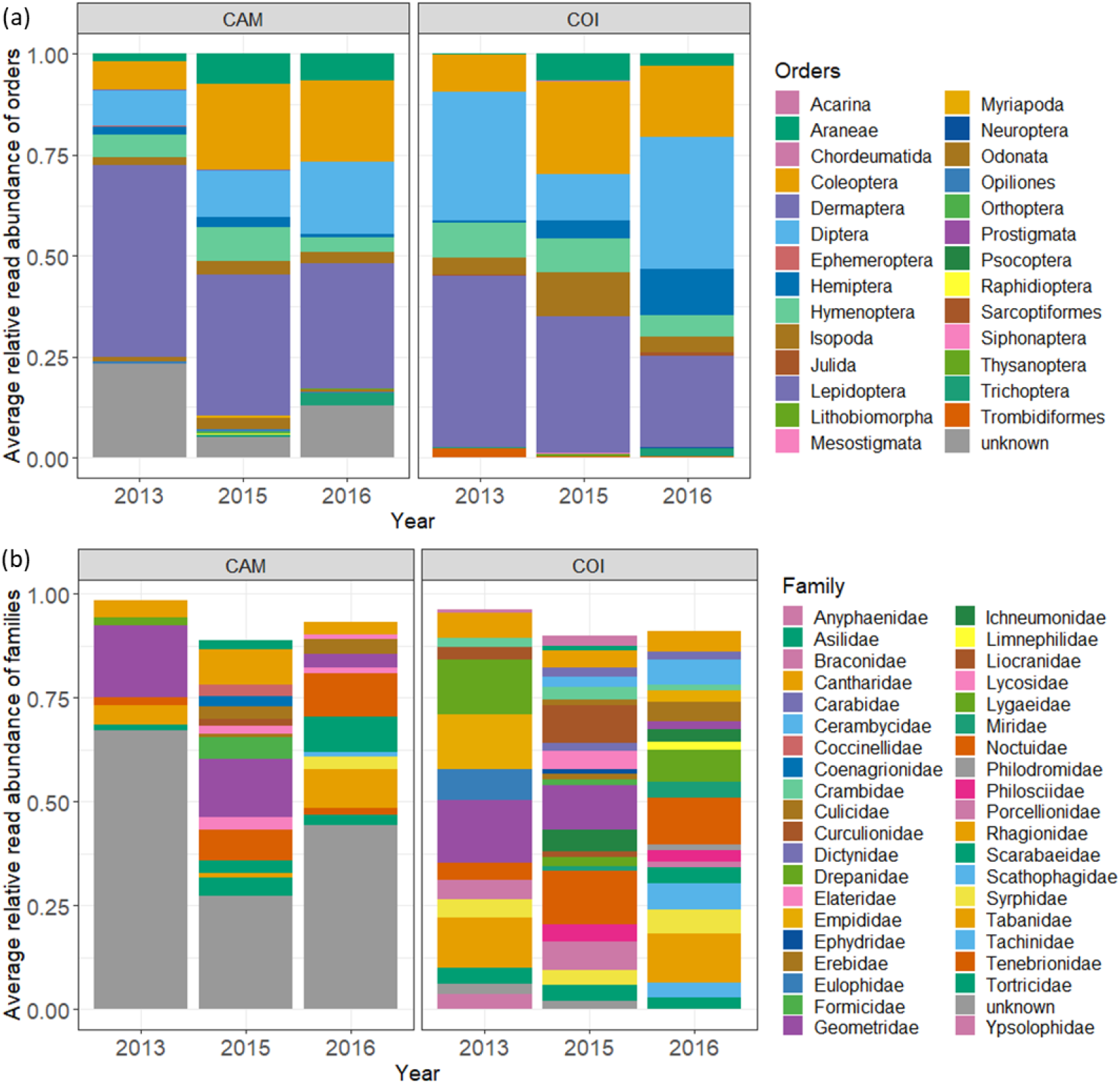
Average relative read abundance (RRA) of arthropod taxa in camera sessions (CAM; where ‘reads’ are size-adjusted prey counts) and in metabarcodes from faeces (COI) collected during camera sessions, visualized as (a) orders and (b) families. The average RRA of an order is the summed proportions of reads over all samples divided by the number of samples.

On the level of order, the FOO of taxa detected in COI correlated less strongly with the relative taxa abundance in size-adjusted prey counts (~biomass) detected on camera (Fig. 5a) (R = 0. 65 (0.33-0.84, p = 0.0007) than the average RRA of taxa in COI (R = 0.85 (0.68-0.94, p < 0.0001; Fig. 5b). The correlations with RRA were consistent across three study years (Fig. 5c): 2013 (R = 0.76 (0.39-0.92, p = 0.001), 2015 (R = 0.92 (0.80-0.97, p < 0.0001) and 2016 (R = 0.78 (0.51-0.91, p < 0.0001). Note that 2013 had a small dataset of five samples. On the family level, the correlation with taxa abundance on camera also was less strong for FOO (R = 0.62 (0.50-0.71, p < 0.0001) than for RRA (R = 0.75 (0.67-0.82, p < 0.0001) (Fig. 5d,e).

**FIGURE 5.**
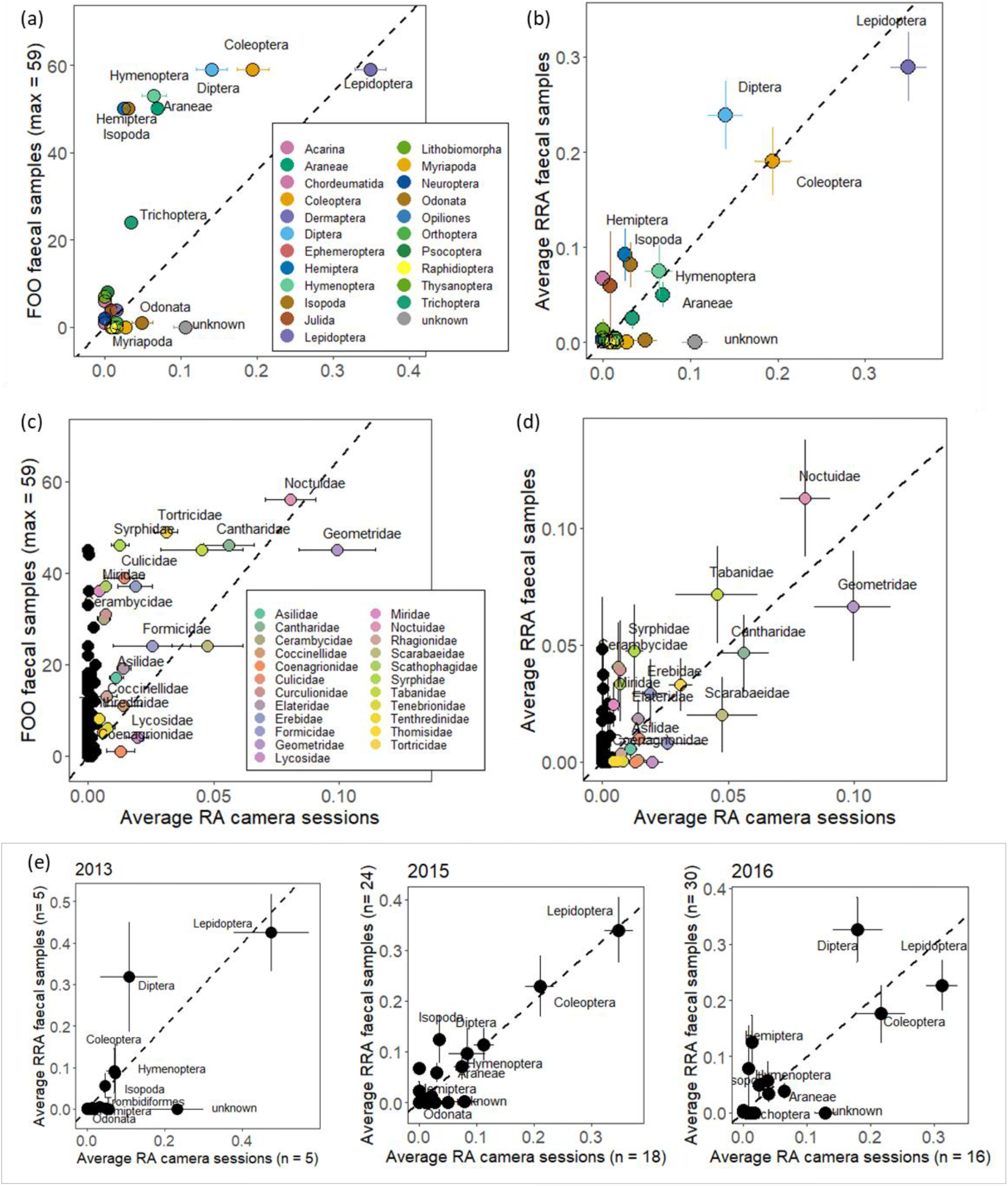
Validation test of retrieval of arthropod taxa through metabarcoding of bird faeces. Frequency of occurrence (FOO) and relative read abundance (RRA) of taxa in faeces collected during camera sessions were determined with COI barcodes. Shown are the relationships of FOO (a, c) and RRA (b, d) of taxa in faecal samples (*N* = 59) versus observed relative prey item (“read”) abundance (RA) of taxa in camera sessions (*N* = 39), for the aggregated study years for orders (a,b) and families (c, d). Also shown are the three separate study years ((e) orders only). The dashed lines illustrate the X = Y relationship to guide the eye.

**In summary, we validated that the relative abundance of consumed taxa can be approximated by the RRA of taxa in faecal samples.**

## 4 DISCUSSION

### 4.1 Proof of principle: what we have learned?

We demonstrated that DNA metabarcoding of faeces with a single COI primer pair can quantitatively retrieve the arthropod diet of an insectivorous bird, where relative read abundance (RRA) of taxa described the observed diet better than the frequency of occurrence (FOO) of taxa.

Our study design, including validation, involved (i) adjusting the DNA-extraction method to increase yield of target COI reads, (ii) adjusting PCR primers to avoid avian DNA, (iii) establishing that low sequencing depths of 2,000-10,000 reads per sample were sufficient, (iv) demonstrating that digestion did not bias taxa recovery, and (v) validating whether diets as estimated with COI metabarcoding corresponded quantitatively with the actual provided diet.

Aggregating data of 5-18 camera sessions per year (n = 3 years), the results of COI metabarcoding matched the camera records on the arthropod order and family level, validating that at these taxonomic levels the relative abundance of taxa could be recovered with metabarcoding. Within the orders Lepidoptera, Diptera and Coleoptera, COI barcodes and camera records detected the same general families. Although both methods were equally good in assessing diet composition of common taxa, metabarcoding had an advantage over camera recordings because with COI barcodes lower taxa were described.

### 4.2 How important is validation?

Validation studies using mock communities or captive animals fed a known diet, have been conducted for a wide range of consumers and prey and have shown that although broad correlations are likely, especially at level of the presence/absence of prey taxa, biases may occur that need to be accounted for (reviewed in Deagle et al. (2019), see also (Thuo et al., 2019). Therefore Deagle et al. (2019) recommended to incorporate cross-validation in a study setup whenever possible.

We found that the approximated biomass of arthropod taxa in the diet could be quantitatively retrieved with deviations within an order of magnitude. This is in contrast with earlier studies using composed arthropod communities (Elbrecht & Leese, 2015; Piñol et al., 2015) which showed that the recovered read abundance per taxon could vary by two to four orders of magnitude from the biomass in the mock community (see also Krehenwinkel et al., 2017; Jusino et al., 2019). In Piñol et al. (2015), especially the read abundance of spiders was underrepresented. Piñol et al. (2015) used the “Zeale” primers developed by Zeale et al. (2011) for the COI locus in arthropods, and pointed out the high number of mismatches between spider template and primers. Underestimation of spiders with the Zeale primers has been reported before (Aldasoro et al., 2019; da Silva et al., 2019). Following King et al., (2015) we used the general invertebrate “Folmer-Brown” primers (Folmer et al., 1994; Brown et al., 2012), which retrieved spiders much better (especially our modified primers, see Step 2). Elbrecht & Leese (2015) used the original Folmer primer pair, and also attributed the discrepancy to mismatches between template and primers. They reported low taxa recovery within Diptera which we did not see in our study with the modified Folmer-Brown primers. Krehenwinkel et al. (2017) used modified Zeale and Folmer primers in various combinations but were unable to amplify some Acari and Hymenoptera. Recently a new COI primer pair with a higher arthropod taxon rate than the Zeale-primers was developed, which had only a slight mismatch between the recovered read abundance and the mock community (ANML primers, Jusino et al., 2019). This highlights that primer choice and PCR is crucial. The reported mismatches also indicate that to retrieve diet on the species level precise calibration is needed (see Krehenwinkel et al. (2017) for guidance).

For a validation study we think that capturing the “real” arthropod prey community is an advantage, but we acknowledge that our camera observations also had biases and that causes of disagreements between camera and COI data can be disputed. Nevertheless, we detected a quantitative match between prey biomass and RRA on the taxonomic levels for which the camera data was reliable: order and (and to a lesser extent) family. We showed that using FOO as metric captured the recorded prey biomass not as well as the RRA. We therefore conclude that our protocol allows for a quantitative use of RRA on the order and family level in insectivorous songbirds, and we think that especially (1) the improved arthropod template-primer match, (2) the removal of uric acids from the DNA template, and (3) the low, forgiving, annealing temperature in triplicate PCRs are important (see also Krehenwinkel et al., 2017). We recommend that future studies on insectivorous birds should test the modified Folmer-Brown primers (this study) and the ANML primers (Jusino et al., 2019) and ideally include a validation with known diet to confirm this protocol indeed works with other types of insectivorous songbirds.

### 4.3 Potential biases - field sample collection and diet assessment

#### Biases introduced by field sample collection

In this validation study a variable temporal gap existed between observed consumption of prey during the camera observations and the collection of faeces for COI analysis. Also, we collected faeces of individual chicks, while we recorded prey brought to the whole brood. By chance the diet of the sampled chick(s) may have deviated from the average brood level diet. An alternative calibration test could have been feeding individual birds specific diets after a period of food deprivation. However, the disadvantage of such detailed calibrations in the case of arthropod diets is that they only include what prey are accessible to the researcher and what individuals want to eat in captivity. Therefore we choose for a rather coarse calibration scheme, using random chicks in a brood and accepting that collected faeces after camera sessions may also reflect what chicks have eaten earlier. We found that COI metabarcoding of faeces could quantitatively retrieve the observed diet, when using aggregated faecal data, as we did with 5-18 camera sessions each year. Only in 2016 two faecal samples per nest were consistently collected right after the camera recording, but this did not improve the match between observed diet and COI data very much (Fig. 5e). This suggests that in this study area diets may have been rather stable within nest (which is supported by diets of multiple chicks within a nest being more similar than between nests), and also across subsequent days (Nicolaus et al., 2019). The observed close quantitative match between prey communities in gizzard and intestine also supports the quantitative use of metabarcoding for determining diets at the level of individuals. Nevertheless, it remains to be tested what period of prey ingestion a single faecal sample represents.

#### Reliability of taxonomic assignment and diversity in camera records

In the camera records, some food items were not taxonomically assigned and various prey taxa detected with COI metabarcoding were not seen on camera (Fig. 4). Clearly small and inconspicuous species were difficult to identify from images, and adults often arrived with a beak filled with multiple prey items obscuring smaller species. Indeed, food items categorized as “unknown” on camera were usually described as small. Especially in 2013 and 2016 the proportion of unassigned food items was relatively high, which may have especially underestimated smaller species (e.g. note the high proportion of Diptera in 2013 and 2016 in COI, Fig. 4). An example of a common family in COI barcodes but not reported on camera very often were the dance flies (Empididae). However, since also larger-sized taxa were sometimes missed on camera, prey size is not the only possible cause of assignment error. Likely, some species resembled other groups, or observers were unfamiliar with specific groups and may have misclassified them, or they sensibly classified species at a higher taxonomic level only. This may explain why at order level, diet composition was very similar between the methods, but at the family level only within Lepidoptera, Diptera and Coleoptera (Fig. 4, Fig S4-8). Especially, taxonomic assignments from camera footage was complicated in Hymenoptera, Hemiptera and Araneae, leading to many “unknowns” (Fig. S4-8) and making quantitative comparisons between datasets within these orders somewhat farfetched.

#### Ecological limitations of COI metabarcoding

General limitations of using COI metabarcoding are that the life stage of prey cannot be assessed (larva versus adult), and ingested items may not have been an intended food item. Indeed parasites such as fleas, mites and parasitoid wasps were common in COI barcodes, and within Hymenoptera, parasitoid wasps (Ichneumonidae, Braconidae and Eulophidae) dominated the COI reads (Fig. S4-8). These wasps could have been ingested through their (caterpillar) hosts, because per camera session only on average four (range 0–9) food items were assigned to Ichneumonidae which amounted to 0.5% overall, and no Braconidae and Eulophidae wasps were reported, and also no fleas or mites. The COI reads contained one order of fleas (Siphonptera), four orders of mites (Mesostigmata, Prostigmata, Sarcoptiformes, Trombidiformes) (Fig. 4). Siphonptera never exceeded a RRA of 0.2%, with on average 0.01% (Fig. 4). Mites, mostly Trombidiformes, had a RRA of 0.4%, but in two samples 7% and 11% (Fig. 4, Fig. S4-5). Braconidae, Eulophidae and Ichneumonidae were represented by 0.1– 2.7% reads. However, this was mostly due to three samples, in which the RRA of Ichneumonid wasps was as high as 97%, 91% and 24%, while being below 0.94% in all other samples (Fig. S4-5).

In summary, parasites in general contributed 1–2% percent to the diet, while Trombidiformes mites and Ichneumonidae wasps were occasionally abundant. We consider it unlikely that these parasites are more sensitive to PCR bias, and therefore think that they were ingested as secondary prey.

### 4.4 Potential biases – technical limitations of COI metabarcoding

#### Taxonomic assignment of COI barcodes

Especially in studies conducted in North America and Western Europe sequencing the COI gene can obtain genus level identification of arthropods using public reference databases such as GenBank and the Barcoding of Life Database BOLD (King et al., 2015). We indeed used a public database for assignment of OTUs, but considered assignments to species level unreliable. Even at the family level some taxa were assigned that do not occur in Europe (e.g. *Phanomorpha*) which again stresses that our validation setup is less precise at family level. Only at the order level, we consider taxonomic assignment error of the retrieved COI reads unlikely. Nevertheless two orders seen on camera as prey items were not detected in COI: Myriapoda (millipedes, centipedes, and others) and Orthoptera (grasshoppers, locusts and crickets). In the case of the myriapods the COI reads were assigned to another millipede order (Chordeumatida). Since we did not taxonomically reassign prey items *a posteriori* this was not corrected in our validation test. Orthoptera were very rarely provided prey items (Fig. 5) and possibly therefore not detected. Orthoptera were detected in faecal samples of adult pied flycatchers using the same methodology (M. Tangili, pers. comm.). For more reliable assignment below order level a reference barcode database of the local prey community is strongly preferred.

#### Taxonomic diversity recovered with COI barcodes

Even though COI barcodes detected a large variety of taxa that matched with recorded diet, the taxonomic diversity of DNA barcodes in a study can potentially be biased (Krehenwinkel et al., 2017; Alberdi et al., 2018) especially when sample size is low. In our validation test, ca. 15% of the faecal samples were dominated by a single species ((Fig. S4-5). This over-dominance was most likely created during laboratory procedures but possibly was not a systematic bias. In Step 2, we established that the two DNA-extraction methods did not show a directional difference in taxa recovery, but when subsampling faeces for DNA-extraction, a single large prey remain may lead to overrepresentation in DNA template. Additionally, PCR can cause overrepresentation of taxa, either randomly by template favouring after the initial PCR cycle, or non-randomly, when primers may match better to some taxa than others. To avoid the risk of PCR bias it is recommended to perform triplicate PCRs on each individual sample before pooling for sequencing (e.g. Vo & Jedlicka, 2014, but see Alberdi et al., 2018). We indeed executed PCR in duplicates or triplicates, and the repeated samples in a triplicate PCR setup in Step 2&3 showed high repeatability of prey composition. To avoid non-random bias, using multiple markers is recommended because it increases the number of taxa recovered (Corse et al., 2019; da Silva et al., 2019), but since this jeopardized testing the relationship between RRA and the observed arthropod taxa to a diet we decided against it. In our protocol the primers did not systematically favour certain taxa (Fig. S4-5) and therefore non-random bias is unlikely.

In conclusion, we propose that the observed over-dominance in some samples may have been caused by transfer of larger fragments in faeces to the DNA-extraction. We reject the hypothesis that over-dominance was an effect of prey digestion, e.g. that degradation of DNA in the digestive track structurally varies between prey species, because we observed very similar prey communities in stomach and intestine content. Dominance of single prey taxa could have been the result of birds consuming large amounts of a single prey species for some time, but that seems unlikely given the camera footage. Remarkably over-dominance by >90% was much more common in the adult males of Step 3 (5 of 16 samples, Fig. S3-2) than in the nestlings of Step 2 (1 of 8 samples) and the validation test (8 of 59 samples, Fig. S4-5). This may hint at different feeding patterns; alternatively the smaller-sized gizzard and intestine samples taken from the males may have been more homogenous. Better insights in the potential causes of over-dominance can come from comparing larger datasets, more detailed calibration feeding experiments, and laboratory study designs with more subsampled duplicates of faeces.

#### Is a single faeces a necessary unit?

Faecal droppings are snap-shots of the diet. In this study we COI barcoded single faeces, which has the advantage that they can be connected to an individual bird. However, single faecal samples can be very limited snap-shots, especially when a dropping is dominated by a single taxa (see examples in Table S4-1). It is cheaper to pool samples if one aims to obtain a general picture of the diet. To assess the effect of pooling samples, we analysed the representation of arthropod orders by averaging individual samples (as presented) and for pooled samples (not shown) which had no significant effect. Although pooling PCR product samples before sequencing may be more cost effective and may not lead to loss of information, it will reduce statistical power, and we will lose important information about individual variation.

### 4.5 Application to ecological studies

A clear added benefit exists of quantifying trophic interactions at the species level, rather than on higher taxonomic levels, as each species has its specific biology, including its niche, phenology, population dynamics, adaptive capacities, etc. Most traditional non-molecular methods to quantify diets of generalist insectivorous organisms are unable to recognize lower taxonomic levels, and hence may miss important ecological interactions. Flycatchers do e.g. feed their nestlings with a rather large fraction of caterpillars (Burger et al., 2012) and the general caterpillar peak (measured by collecting their faeces) has advanced in response to climate change (Both, van Asch, Bijlsma, van den Burg, & Visser, 2009). However, little is known whether all caterpillar species do advance their phenology to a similar extent, and how flycatchers may switch from one caterpillar species to the other depending on its abundance and phenology. If we want to understand how food-webs respond to climate or other environmental changes including its eco-evolutionary dynamics, we need to quantify interactions preferably on the species level, because population and evolutionary dynamics are species characteristics. The accuracy of metabarcoding techniques to identify trophic interactions at the species level may not yet be absolute, but often much better than traditional methods. Furthermore, it allows to study interactions beyond the easily observed nestling periods, and e.g. we may now quantify year-round diet as long as we can obtain faeces of the species of interest.

Metabarcoding may hence be a quantitative tool to study the relative contribution of different prey taxa in the diet of consumers, but since individuals do not survive on relative contributions but rather need absolute amounts of food, it is still necessary to measure intake rates to study how diet composition changes depending on ecological circumstances. Hence we see this method as an important addition to the toolkit of field ecologists, enabling them also to focus more on sampling the temporal and spatial abundance of the prey that really are important for the predator species of interest.

## 5 CONCLUSIONS

The successful validation of our metabarcoding approach opens the opportunity to quantitatively monitor diet and trophic interactions in songbird study systems. Essential technical elements of our approach are (1) host-avoiding, non-degenerative primers, (2) extraction methods avoiding uric acids, (3) low annealing temperatures and triplicate PCRs for high taxonomic resolution, and (4) sequencing library preparation without normalization. However, before application in other study systems we recommend local validation with recorded diets. The taxonomic level of validation can be improved by involving specific taxa experts when analysing camera footage, or by feeding experiments or taxon-specific PCR to establish correction factors for certain prey groups (Zeale et al., 2011; Thomas et al., 2016).

## ACKNOWLEDGEMENTS

Janne Ouwehand collected and donated two precious Africa samples for the explorative initial tests. Rob Bijlsma directed us to the adults used in the digestive bias test. Marco van der Velde gave crucial advice at several stages of the project, which largely improved lab protocols and data handling. Also Pieter van Veelen had helpful suggestions for lab protocols, and very kindly shared his R scripts. Also Marianthi Tangili advised us on R data handling. Kristiaan van der Gaag and Rick de Leeuw took excellent care of library preparation and sequencing. We would like to thank the Center for Information Technology of the University of Groningen for their support and for providing access to the Peregrine high performance computing cluster, and for their invaluable help with the bioinformatics pipeline design and *big data* handling. This research was performed under legal permission of the Ethical Committee for Animal Experimentation of the University of Groningen (DEC 6812). The study was supported by Netherlands Organization for Scientific Research (NWO-ALW, grant number LWOP.2014.109 to MN and CB) and a start-up grant from the GUF-Gratama foundation (grant number 2016-05 to CB, YIV and MD).

## AUTHORS’ CONTRIBUTIONS

The study was designed by CB, YIV and PdK. The photo camera recordings were conducted by RU, MN and JMS. Faecal samples were collected by CB, RU, MN, JMS, MMD and YIV. Photo footage was curated by RU and image analyses were conducted by JMS and CB. The molecular methods were developed by YIV, PdK, MMD and AG. DNA labwork was conducted by YIV and AG. MiSeq sequencing was conducted by PdK. The bioinformatics pipeline and scripts was designed by YIV, MMD, AG and KK. All authors commented on earlier drafts of the manuscript.

## DATA ACCESSIBILITY

1. Raw data: Fastq files - archived at University of Groningen
2. Data tables: OTUtable, TAXAtable, SAMPLE-INFO - Submission docs bioRxiv 2020
3. Scripts: USearch jobscript, Rgui-scripts - Submission docs bioRxiv 2020

## ONLINE SUPPORTING MATERIAL

Additional figures: 13 figures, 6 tables in 4 Supplements

## REFERENCES

Alberdi, A., Aizpurua, O., Bohmann, K., Gopalakrishnan, S., Lynggaard, C., Nielsen, M., & Gilbert, M. T. P. (2019). Promises and pitfalls of using high-throughput sequencing for diet analysis. Molecular Ecology Resources, 19(2), 327–348. doi:10.1111/1755-0998.12960

Alberdi, A., Aizpurua, O., Gilbert, M. T. P., & Bohmann, K. (2018). Scrutinizing key steps for reliable metabarcoding of environmental samples. Methods in Ecology and Evolution, 9(1), 134–147. doi:10.1111/2041-210X.12849

Aldasoro, M., Garin, I., Vallejo, N., Baroja, U., Arrizabalaga-Escudero, A., Goiti, U., & Aihartza, J. (2019). Gaining ecological insight on dietary allocation among horseshoe bats through molecular primer combination. PLoS ONE, 14(7), 1–15. doi:10.1371/journal.pone.0220081

Anderson, M. J. (2001). A new method for non-parametric multivariate analysis of variance. Austral Ecology, 26(1), 32–46. doi:10.1046/j.1442-9993.2001.01070.x

Ando, H., Setsuko, S., Horikoshi, K., Suzuki, H., Umehara, S., Inoue-Murayama, M., & Isagi, Y. (2013). Diet analysis by next-generation sequencing indicates the frequent consumption of introduced plants by the critically endangered red-headed wood pigeon (*Columba janthina nitens*) in oceanic island habitats. Ecology and Evolution, 3(12), 4057–4069. doi:10.1002/ece3.773

Benson, D. A., Karsch-mizrachi, I., Lipman, D. J., Ostell, J., & Sayers, E. W. (2009). GenBank. Nucleic Acids Research, 37, D26–D31. doi:10.1093/nar/gkn723

Both, C., Bijlsma, R. G., & Ouwehand, J. (2016). Repeatability in spring arrival dates in Pied Flycatchers varies among years and sexes. Ardea, 104(1), 3–21. doi:10.5253/arde.v104i1.a1

Both, C., Bouwhuis, S., Lessells, C. M., & Visser, M. E. (2006). Climate change and population declines in a long-distance migratory bird. Nature, 441(1), 81–83. doi:10.1038/nature04539

Both, C., van Asch, M., Bijlsma, R. G., van den Burg, A. B., & Visser, M. E. (2009). Climate change and unequal phenological changes across four trophic levels: Constraints or adaptations? Journal of Animal Ecology, 78(1), 73–83. doi:10.1111/j.1365-2656.2008.01458.x

Brandon-Mong, G. J., Gan, H. M., Sing, K. W., Lee, P. S., Lim, P. E., & Wilson, J. J. (2015). DNA metabarcoding of insects and allies: An evaluation of primers and pipelines. Bulletin of Entomological Research, 105(6), 717–727. doi:10.1017/S0007485315000681

Brown, D. S., Jarman, S. N., & Symondson, W. O. C. (2012). Pyrosequencing of prey DNA in reptile faeces: Analysis of earthworm consumption by slow worms. Molecular Ecology Resources, 12(2), 259–266. doi:10.1111/j.1755-0998.2011.03098.x

Burger, C., Belskii, E., Eeva, T., Laaksonen, T., Mägi, M., Mänd, R., … Both, C. (2012). Climate change, breeding date and nestling diet: How temperature differentially affects seasonal changes in pied flycatcher diet depending on habitat variation. Journal of Animal Ecology, 81(4), 926–936. doi:10.1111/j.1365-2656.2012.01968.x

Cholewa, M., & Wesołowski, T. (2011). Nestling Food of European Hole-Nesting Passerines: Do We Know Enough to Test the Adaptive Hypotheses on Breeding Seasons? Acta Ornithologica, 46(2), 105–116. doi:10.3161/000164511x625874

Corse, E., Tougard, C., Archambaud-Suard, G., Agnèse, J. F., Messu Mandeng, F. D., Bilong Bilong, C. F., … Dubut, V. (2019). One-locus-several-primers: A strategy to improve the taxonomic and haplotypic coverage in diet metabarcoding studies. Ecology and Evolution, 9(8), 4603–4620. doi:10.1002/ece3.5063

da Silva, L. P., Mata, V. A., Lopes, P. B., Pereira, P., Jarman, S. N., Lopes, R. J., & Beja, P. (2019). Advancing the integration of multi-marker metabarcoding data in dietary analysis of trophic generalists. Molecular Ecology Resources, 19(6), 1420–1432. doi:10.1111/1755-0998.13060

Deagle, B. E., Thomas, A. C., McInnes, J. C., Clarke, L. J., Vesterinen, E. J., Clare, E. L., … Eveson, J. P. (2019). Counting with DNA in metabarcoding studies: How should we convert sequence reads to dietary data? Molecular Ecology, 28(2), 391–406. doi:10.1111/mec.14734

Dixon, P. (2003). VEGAN, a package of R functions for community ecology. Journal of Vegetation Science, 14, 927–930. Retrieved from http://doi.wiley.com/10.1111/j.1654-1103.2002.tb02049.x

Edgar, R. C. (2010). Search and clustering orders of magnitude faster than BLAST. Bioinformatics, 26(19), 2460–2461. doi:10.1093/bioinformatics/btq461

Elbrecht, V., & Leese, F. (2015). Can DNA-based ecosystem assessments quantify species abundance? Testing primer bias and biomass-sequence relationships with an innovative metabarcoding protocol. PLoS ONE, 10(7), 1–16. doi:10.1371/journal.pone.0130324

Elton, C. S. (1927). Chapter V: The animal community. In Animal ecology, by Charles Elton; with an introduction by Julian S. Huxley. (pp. 1–15). doi:10.5962/bhl.title.7435

Evans, D. M., Kitson, J. J. N., Lunt, D. H., Straw, N. A., & Pocock, M. J. O. (2016). Merging DNA metabarcoding and ecological network analysis to understand and build resilient terrestrial ecosystems. Functional Ecology, 30(12), 1904–1916. doi:10.1111/1365-2435.12659

Folmer, O., Black, M., Hoeh, W., Lutz, R., & Vrijenhoek, R. (1994). DNA primers for amplification of mitochondrial cytochrome c oxidase subunit I from diverse metazoan invertebrates. Molecular Marine Biology and Biotechnology, 3(5), 294–299.

Fox, J., Friendly, M., & Weisberg, S. (2013). Hypothesis tests for multivariate linear models using the car package. R Journal, 5(1), 39–52. doi:10.32614/rj-2013-004

Gerwing, T. G., Kim, J.-H., Hamilton, D. J., Barbeau, M. A., & Addison, J. A. (2016). Diet reconstruction using next-generation sequencing increases the known ecosystem usage by a shorebird. The Auk, 133(2), 168–177. doi:10.1642/auk-15-176.1

Hallmann, C. A., Foppen, R. P. B., van Turnhout, C. A. M., de Kroon, H., & Jongejans, E. (2014). Declines in insectivorous birds are associated with high neonicotinoid concentrations. Nature, 511(7509), 341–343. doi:10.1038/nature13531

Hebert, P. D. N., Ratnasingham, S., & DeWaard, J. R. (2003). Barcoding animal life: Cytochrome c oxidase subunit 1 divergences among closely related species. Proceedings of the Royal Society B: Biological Sciences, 270(SUPPL. 1), 96–99. doi:10.1098/rsbl.2003.0025

Ishii, K., & Fukui, M. (2001). Optimization of Annealing Temperature To Reduce Bias Caused by a Primer Mismatch in Multitemplate PCR. Applied and Environmental Micriobiology, 67(8), 3753–3755. doi:10.1128/AEM.67.8.3753

Jedlicka, J. A., Sharma, A. M., & Almeida, R. P. P. (2013). Molecular tools reveal diets of insectivorous birds from predator fecal matter. Conservation Genetics Resources, 5(3), 879–885. doi:10.1007/s12686-013-9900-1

Jusino, M. A., Banik, M. T., Palmer, J. M., Wray, A. K., Xiao, L., Pelton, E., … Lindner, D. L. (2019). An improved method for utilizing high-throughput amplicon sequencing to determine the diets of insectivorous animals. Molecular Ecology Resources, 19(1), 176–190. doi:10.1111/1755-0998.12951

Kearse, M., Moir, R., Wilson, A., Stones-Havas, S., Cheung, M., Sturrock, S., … Drummond, A. (2012). Geneious Basic: An integrated and extendable desktop software platform for the organization and analysis of sequence data. Bioinformatics, 28(12), 1647–1649. doi:10.1093/bioinformatics/bts199

King, R. A., Read, D. S., Traugott, M., & Symondson, W. O. C. (2008). Molecular analysis of predation: A review of best practice for DNA-based approaches. Molecular Ecology, 17(4), 947–963. doi:10.1111/j.1365-294X.2007.03613.x

King, R. A., Symondson, W. O. C., & Thomas, R. J. (2015). Molecular analysis of faecal samples from birds to identify potential crop pests and useful biocontrol agents in natural areas. Bulletin of Entomological Research, 105(3), 261–272. doi:10.1017/S0007485314000935

Krehenwinkel, H., Kennedy, S. R., Rueda, A., Lam, A., & Gillespie, R. G. (2018). Scaling up DNA barcoding – Primer sets for simple and cost efficient arthropod systematics by multiplex PCR and Illumina amplicon sequencing. Methods in Ecology and Evolution, 9(11), 2181–2193. doi:10.1111/2041-210X.13064

Krehenwinkel, H., Wolf, M., Lim, J. Y., Rominger, A. J., Simison, W. B., & Gillespie, R. G. (2017). Estimating and mitigating amplification bias in qualitative and quantitative arthropod metabarcoding. Scientific Reports, 7(1), 1–12. doi:10.1038/s41598-017-17333-x

Kress, W. J., García-Robledo, C., Uriarte, M., & Erickson, D. L. (2015). DNA barcodes for ecology, evolution, and conservation. Trends in Ecology and Evolution, 30(1), 25–35. doi:10.1016/j.tree.2014.10.008

Krüger, F., Clare, E. L., Symondson, W. O. C., Keišs, O., & Petersons, G. (2014). Diet of the insectivorous bat *Pipistrellus nathusii* during autumn migration and summer residence. Molecular Ecology, 23(15), 3672–3683. doi:10.1111/mec.12547

McClenaghan, B., Nol, E., & Kerr, K. C. R. (2019). DNA metabarcoding reveals the broad and flexible diet of a declining aerial insectivore. Auk, 136(1), 1–11. doi:10.1093/auk/uky003

McMurdie, P. J., & Holmes, S. (2013). Phyloseq: An R Package for Reproducible Interactive Analysis and Graphics of Microbiome Census Data. PLoS ONE, 8(4). doi:10.1371/journal.pone.0061217

Merilä, J., & Wiggins, D. A. (1995). Interspecific Competition for Nest Holes Causes Adult Mortality in the Collared Flycatcher. The Condor, 97(2), 445–450. Retrieved from https://www.jstor.org/stable/1369030

Nicolaus, M., Barrault, S. C. Y., & Both, C. (2019). Diet and provisioning rate differ predictably between dispersing and philopatric pied flycatchers. Behavioral Ecology, 30(1), 114–124. doi:10.1093/beheco/ary152

Oksanen, J., Blanchet, F. G., Friendly, M., Kindt, R., Legendre, P., Mcglinn, D., … Wagner, H. (2019). vegan: Community Ecology Package. R package version 2.4-2. Community Ecology Package. Retrieved from https://cran.r-project.org/web/packages/vegan/vegan.pdf

Piñol, J., Mir, G., Gomez-Polo, P., & Agustí, N. (2015). Universal and blocking primer mismatches limit the use of high-throughput DNA sequencing for the quantitative metabarcoding of arthropods. Molecular Ecology Resources, 15(4), 819–830. doi:10.1111/1755-0998.12355

Piñol, J., San Andrés, V., Clare, E. L., Mir, G., & Symondson, W. O. C. (2014). A pragmatic approach to the analysis of diets of generalist predators: The use of next-generation sequencing with no blocking probes. Molecular Ecology Resources, 14(1), 18–26. doi:10.1111/1755-0998.12156

Pompanon, F., Deagle, B. E., Symondson, W. O. C., Brown, D. S., Jarman, S. N., & Taberlet, P. (2012). Who is eating what: Diet assessment using next generation sequencing. Molecular Ecology, 21(8), 1931–1950. doi:10.1111/j.1365-294X.2011.05403.x

Ratnasingham, S., & Hebert, P. D. (2007). BOLD: The Barcode of Life Data System (http://www.boldsystems.org). Molecular Ecology Notes, 7(3), 355–364. doi:10.1111/j.1471-8286.2006.01678.x

Rytkönen, S., Vesterinen, E. J., Westerduin, C., Leviäkangas, T., Vatka, E., Mutanen, M., … Orell, M. (2019). From feces to data: A metabarcoding method for analyzing consumed and available prey in a bird-insect food web. Ecology and Evolution, 9(1), 631–639. doi:10.1002/ece3.4787

Samplonius, J. M., & Both, C. (2019). Climate Change May Affect Fatal Competition between Two Bird Species. Current Biology, 29(2), 327–331.e2. doi:10.1016/j.cub.2018.11.063

Samplonius, J. M., Kappers, E. F., Brands, S., & Both, C. (2016). Phenological mismatch and ontogenetic diet shifts interactively affect offspring condition in a passerine. Journal of Animal Ecology, 85(5), 1255–1264. doi:10.1111/1365-2656.12554

Shutt, J. D., Nicholls, J. A., Trivedi, U. H., Burgess, M. D., Stone, G. N., Hadfield, J. D., & Phillimore, A. B. (2020). Gradients in richness and turnover of a forest passerine’s diet prior to breeding: A mixed model approach applied to faecal metabarcoding data. Molecular Ecology, 29(6), 1199–1213. doi:10.1111/mec.15394

Slagsvold, T. (1975). Competition between the Great Tit *Parus major* and the Pied Flycatcher *Ficedula hypoleuca* in the Breeding Season. Ornis Scandinavica, 6(2), 179–190. Retrieved from https://www.jstor.org/stable/3676230

Taberlet, P., Bonin, A., Zinger, L., & Coissac, E. (2018). Environmental DNA: For Biodiversity Research and Monitoring. Oxford University Press. doi:10.1093/oso/9780198767220.001.0001

Thomas, A. C., Deagle, B. E., Eveson, J. P., Harsch, C. H., & Trites, A. W. (2016). Quantitative DNA metabarcoding: Improved estimates of species proportional biomass using correction factors derived from control material. Molecular Ecology Resources, 16(3), 714–726. doi:10.1111/1755-0998.12490

Thuo, D., Furlan, E., Broekhuis, F., Kamau, J., Macdonald, K., & Gleeson, D. M. (2019). Food from faeces: Evaluating the efficacy of scat DNA metabarcoding in dietary analyses. PLoS ONE, 14(12), 1–15. doi:10.1371/journal.pone.0225805

Trevelline, B. K., Nuttle, T., Hoenig, B. D., Brouwer, N. L., Porter, B. A., & Latta, S. C. (2018). DNA metabarcoding of nestling feces reveals provisioning of aquatic prey and resource partitioning among Neotropical migratory songbirds in a riparian habitat. Oecologia, 187(1), 85–98. doi:10.1007/s00442-018-4136-0

Valentini, A., Pompanon, F., & Taberlet, P. (2009). DNA barcoding for ecologists. Trends in Ecology and Evolution, 24(2), 110–117. doi:10.1016/j.tree.2008.09.011

Vo, A. T. E., & Jedlicka, J. A. (2014). Protocols for metagenomic DNA extraction and Illumina amplicon library preparation for faecal and swab samples. Molecular Ecology Resources, 14(6), 1183–1197. doi:10.1111/1755-0998.12269

Ward, R. D. (2009). DNA barcode divergence among species and genera of birds and fishes. Molecular Ecology Resources, 9(4), 1077–1085. doi:10.1111/j.1755-0998.2009.02541.x

Wirta, H. K., Vesterinen, E. J., Hambäck, P. A., Weingartner, E., Rasmussen, C., Reneerkens, J., … Roslin, T. (2015). Exposing the structure of an Arctic food web. Ecology and Evolution, 5(17), 3842–3856. doi:10.1002/ece3.1647

Wong, C. K., Chiu, M. C., Sun, Y. H., Hong, S. Y., & Kuo, M. H. (2015). Using molecular scatology to identify aquatic and terrestrial prey in the diet of a riparian predator, the Plumbeous Water Redstart *Phoenicurus fuliginosa*. Bird Study, 62(3), 368–376. doi:10.1080/00063657.2015.1032888

Zeale, M. R. K., Butlin, R. K., Barker, G. L. A., Lees, D. C., & Jones, G. (2011). Taxon-specific PCR for DNA barcoding arthropod prey in bat faeces. Molecular Ecology Resources, 11(2), 236–244. doi:10.1111/j.1755-0998.2010.02920.x

