## Supplemental Figures and Tables for "DNA metabarcoding successfully quantifies relative abundances of arthropod taxa in songbird diets: a validation study using camera-recorded diets"

To:

### Supplement 1 of 4

#### Step 1: retrieval of target Phylum Arthropoda

We made an initial evaluation of published primers to target Arthropoda through COI metabarcoding of faecal samples. Here additional information is given about the yield per taxa (**Fig. S1-1**) using the generic invertebrate primers generic invertebrate COI primers LCO1490 (Folmer, Black, Hoeh, Lutz, & Vrijenhoek, 1994) and HCO1777 (Brown, Jarman, & Symondson, 2012).

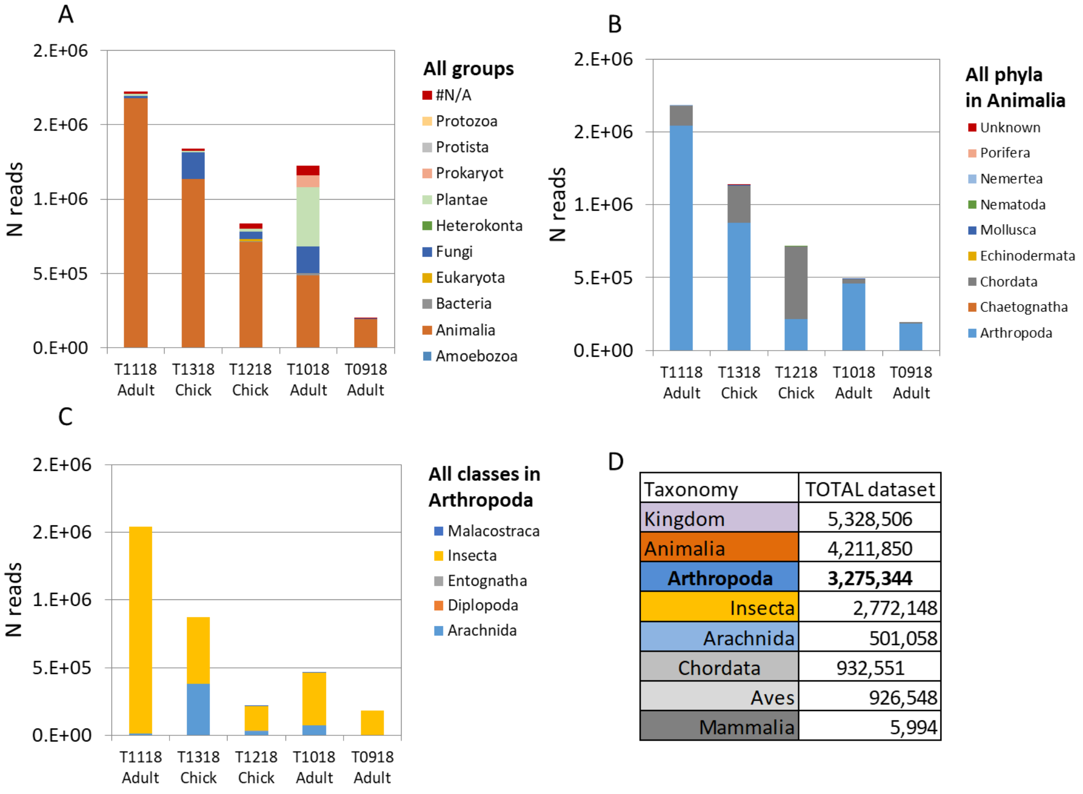


| Taxonomy | TOTAL dataset |
|---|---|
| Kingdom | 5,328,506 |
| Animalia | 4,211,850 |
| **Arthropoda** | **3,275,344** |
| Insecta | 2,772,148 |
| Arachnida | 501,058 |
| Chordata | 932,551 |
| Aves | 926,548 |
| Mammalia | 5,994 |

**FIGURE S1-1** Number of reads in observed taxa in five faecal samples: three samples taken in The Netherlands (T1118, T1318, T1218) and two samples taken in Africa (T1018, T0198). (A) Numbers of reads assigned to Animalia and to non-target groups such as plants and fungi; #N/A is the category of reads that could not be assigned. (B) Number of reads for two Animalia phyla: Arthropoda and Chordata. The remaining detected phyla are listed in the legend, but too rare to be visible in the bar graph. The proportion Unknown indicates reads assigned to OTUs that were not assigned to a taxa. (C) Contribution of the five classes detected in target taxa Arthropoda: some Malacostraca, Entognatha, Diplopod, but mostly Insecta and Arachnida. (D) Overview of numbers shown in bar graphs. Note that most Chordata reads were assigned to Aves.

#### Supplement 2 of 4

##### Step 2: primer redesign, DNA extraction method and read depth

This experiment is an evaluation of (1) DNA extraction methods, (2) modified COI primers, and (3) read/sequencing depth.

In this supplement additional information is given about the PCR primer sequence (**Table S2-1**) taxa used for primer re-design (**Table S2-2**), pairwise comparison of RRA per arthropod order (**Table S2-3**), the methodological evaluation of primers and DNA extraction (**Fig. S2-1**) and sequencing depth in a short and extended sequencing run (**Fig. S2-2**).

**TABLE S2-1** Overview of the original and modified COI primers and the corresponding primer sequences. The variations are indicated in bold. Note that nomenclature refers to the base on the locus.

| COI primer | Forward | Sequence | Source |
|---|---|---|---|
| **Original** | LCO1490 | 5'-GGTCAACAAATCATAAAGATATTGG-3' | Folmer et al. 1994 |
| **Modified** | LCO1490_5T | 5'-GGTC**T**ACAAATCATAAAGATATTGG-3' | this study |
| | **Reverse** | | |
| **Original** | HCO1777 | 5'-ACTTATATTGTTTATACGAGGGAA-3' | Brown et al. 2012 |
| **Modified** | HCO1777_15T | 5'-ACTTATATT**A**TTTATACGAGGGAA-3' | this study |
| **Modified** | HCO1777_15T_24A | 5'-**T**CTTATATT**A**TTTATACGAGGGAA-3' | this study |

**TABLE S2-2** Overview of taxa and GenBank accession numbers used to design primers that are more Arthropod specific than the published primers given in Table S2-1.

| Species | Order | Accession number GenBank |
|---|---|---|
| *Cyclosa conica* | Araneae | KP655120 |
| *Athous subfucus* | Coleoptera | KF317270 |
| *Cantharis pellucida* | Coleoptera | KM447512 |
| *Empis bicuspidata* | Diptera | KF297873 |
| *Scathophaga furcata* | Diptera | KP037602 |
| *Psallus varians* | Hemiptera | KM021773 |
| *Deraeocoris lutescens* | Hemiptera | KJ541653 |
| *Miris striatus* | Hemiptera | KM022933 |
| *Formica sanguinea* | Hymenoptera | KX665113 |
| *Lasius niger* | Hymenoptera | AB007981 |
| *Apocheima hispidaria* | Lepidoptera | GU654879 |
| *Polyploca ridens* | Lepidoptera | JF860168 |
| *Ditula angustiorana* | Lepidoptera | FJ412377 |
| *Ficedula hypoleuca* | Passeriformes | GU571891 |

**TABLE S2-3** Pairwise comparison of RRA per arthropod order in Step 2. This experiment compared two DNA extraction methods, and original versus modified primers. Given are RRA per order. S1 and S2 refer to sample IDs. Extraction methods were PowerSoil (Qiagen DNeasy *PowerSoil* Kit) and PureLink (Invitrogen™ *PureLink*™ Microbiome DNA Purification Kit). Each DNA extraction was tested with the original and modified primers. Original primers: LCO1490 and HCO1777; modified primers: LCO1490_5T (5'-GGTCTACAAATCATAAAGATATTGG-3') and HCO1777_15T (5'-ACTTATATTATTTATACGAGGGAA-3').

| Arthropod orders | original | modified | original | modified | original | modified | original | modified |
|---|---|---|---|---|---|---|---|---|
| Araneae | 3.01% | 8.46% | 1.82% | 7.98% | 2.09% | 5.99% | 0.03% | 0.14% |
| Coleoptera | 0.92% | 0.08% | 0.81% | 0.58% | 46.11% | 23.28% | 1.73% | 0.36% |
| Collembola | 0.51% | 2.10% | 0.55% | 1.71% | 0.00% | 0.00% | 0.00% | 0.00% |
| Diplostraca | 0.00% | 0.00% | 0.00% | 0.00% | 0.00% | 0.00% | 1.11% | 0.00% |
| Diptera | 2.08% | 3.71% | 7.56% | 5.14% | 2.95% | 8.09% | 0.84% | 0.20% |
| Hemiptera | 53.74% | 47.24% | 64.20% | 58.17% | 2.38% | 1.60% | 1.11% | 0.11% |
| Hymenoptera | 11.47% | 13.33% | 8.91% | 10.43% | 37.43% | 55.46% | 90.35% | 98.96% |
| Isopoda | 4.98% | 3.88% | 5.85% | 3.37% | 0.00% | 0.00% | 0.00% | 0.00% |
| Lepidoptera | 21.81% | 19.41% | 9.53% | 12.40% | 0.02% | 0.11% | 0.40% | 0.08% |
| Mesostigmata | 1.43% | 1.40% | 0.55% | 0.04% | 0.60% | 0.00% | 4.31% | 0.01% |
| Neuroptera | 0.00% | 0.00% | 0.00% | 0.00% | 0.00% | 0.00% | 0.00% | 0.00% |
| Orthoptera | 0.00% | 0.00% | 0.00% | 0.00% | 0.00% | 0.05% | 0.00% | 0.00% |
| Prostigmata | 0.01% | 0.00% | 0.00% | 0.00% | 6.57% | 4.93% | 0.06% | 0.13% |
| Psocoptera | 0.01% | 0.40% | 0.09% | 0.14% | 0.00% | 0.00% | 0.00% | 0.00% |
| Thysanoptera | 0.03% | 0.00% | 0.12% | 0.05% | 0.00% | 0.00% | 0.00% | 0.00% |
| Trichoptera | 0.00% | 0.00% | 0.00% | 0.00% | 0.00% | 0.00% | 0.00% | 0.00% |
| Trombidiformes | 0.00% | 0.00% | 0.00% | 0.00% | 1.86% | 0.49% | 0.06% | 0.01% |
| | **S1 PowerSoil** | | **S1 PureLink** | | **S2 PowerSoil** | | **S2 Purelink** | |

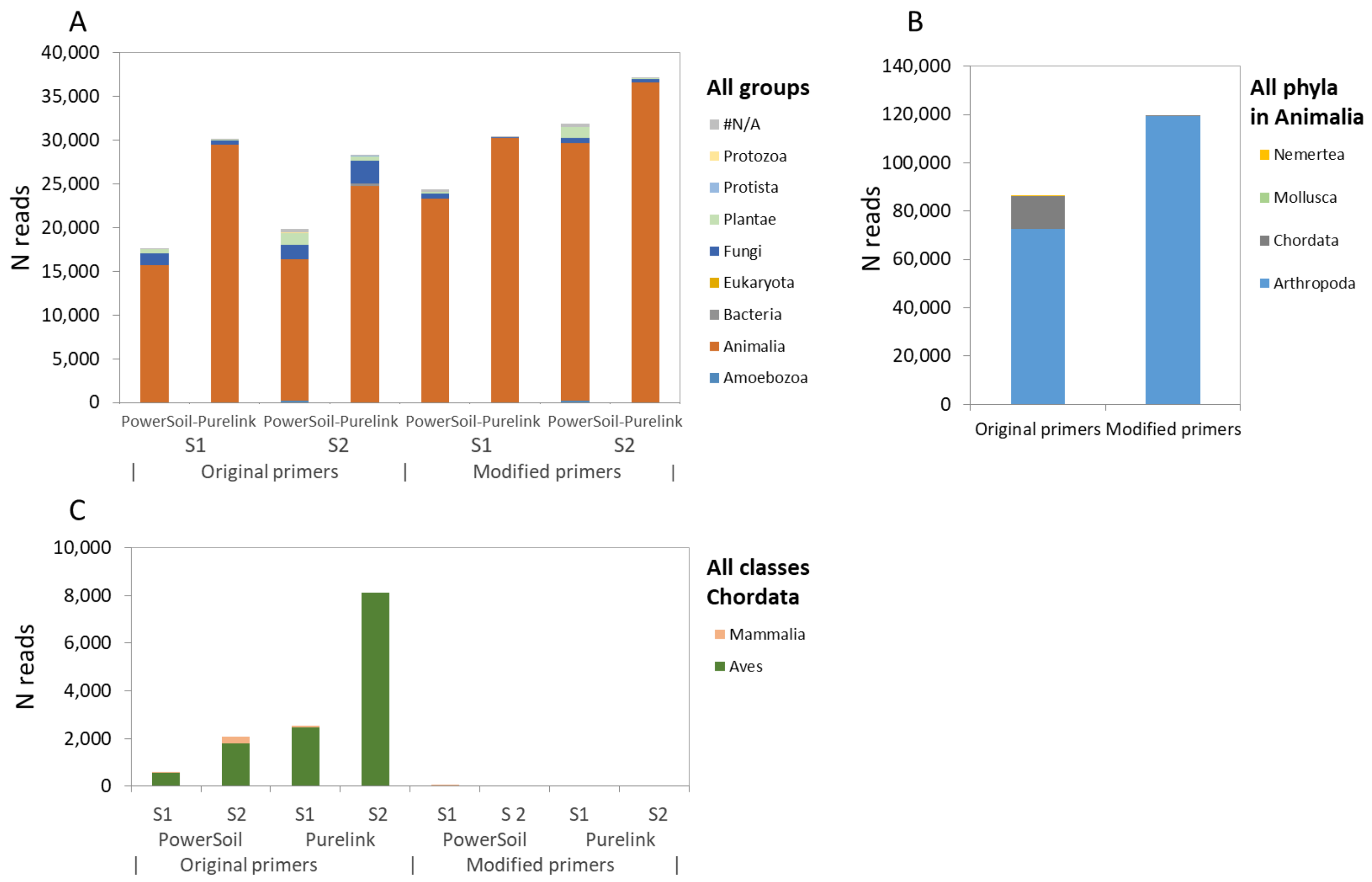


**FIGURE S2-1** Results of methodological evaluation of primers, DNA extraction and sequencing depth. Tested extraction methods were PowerSoil (Qiagen DNeasy *PowerSoil* Kit) and PureLink (Invitrogen™ *PureLink*™ Microbiome DNA Purification Kit). Each DNA extraction was tested with the original and modified primers. Original primers: LCO1490 (Folmer et al., 1994) and HCO1777 (Brown et al., 2012); modified primers: LCO1490_5T (5'-GGTCTACAAATCATAAAGATATTGG-3') and HCO1777_15T (5'-ACTTATATTATTTATACGAGGGAA-3'), this study. (A) Number of reads in all groups observed in four paired tests comparing primers (original primers versus modified primers, see Table S2-1) and DNA extraction methods (PowerSoil versus PureLink) for two samples (S1 and S2). (B) Comparison of primer sets in the detection rate of phyla within Animalia, to test for the occurrence of target Arthropoda and non-target Chordata. (C) Comparison of detection of Aves with Chordata between primer sets and DNA extraction methods, to test whether we successfully reduced the amplification of host DNA.

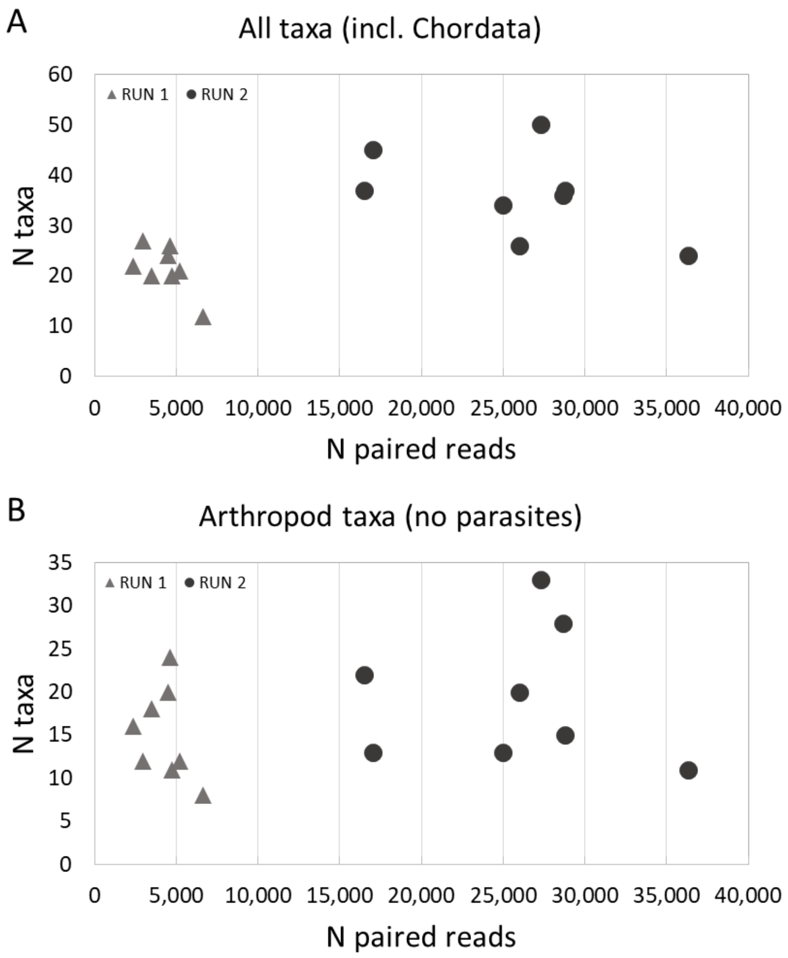


**FIGURE S2-2** Rarefaction plots of number of total taxa and arthropod taxa found against sequencing depth in (A) the limited sequencing run: RUN 1, approximately 10,000 raw sequences per sample and 2,400-6,700 paired reads, and (B) the extended sequencing run: RUN 2, >50,000 sequences per sample yielding 16,500-36,300 paired reads. All taxa includes Chordata, and thus includes reads from the host species. From the arthropod taxa parasites were excluded to assess the effect of read depth on food taxa.

#### Supplement 3 of 4

#### Step 3: digestive bias

Evaluation of effect of digestion on detection of taxa through COI metabarcoding.

In this supplement additional information is given about the detected diversity in gizzard and intestine samples (**Fig. S3-1**) and taxa found in the negative controls (**Table S3-1**). Also shown are the occurrence and (relative) read abundance of taxa detected in both organs (phyla: **Fig. S3-1**; orders: **Fig. S3-2;** genus**: Table S3-2**).

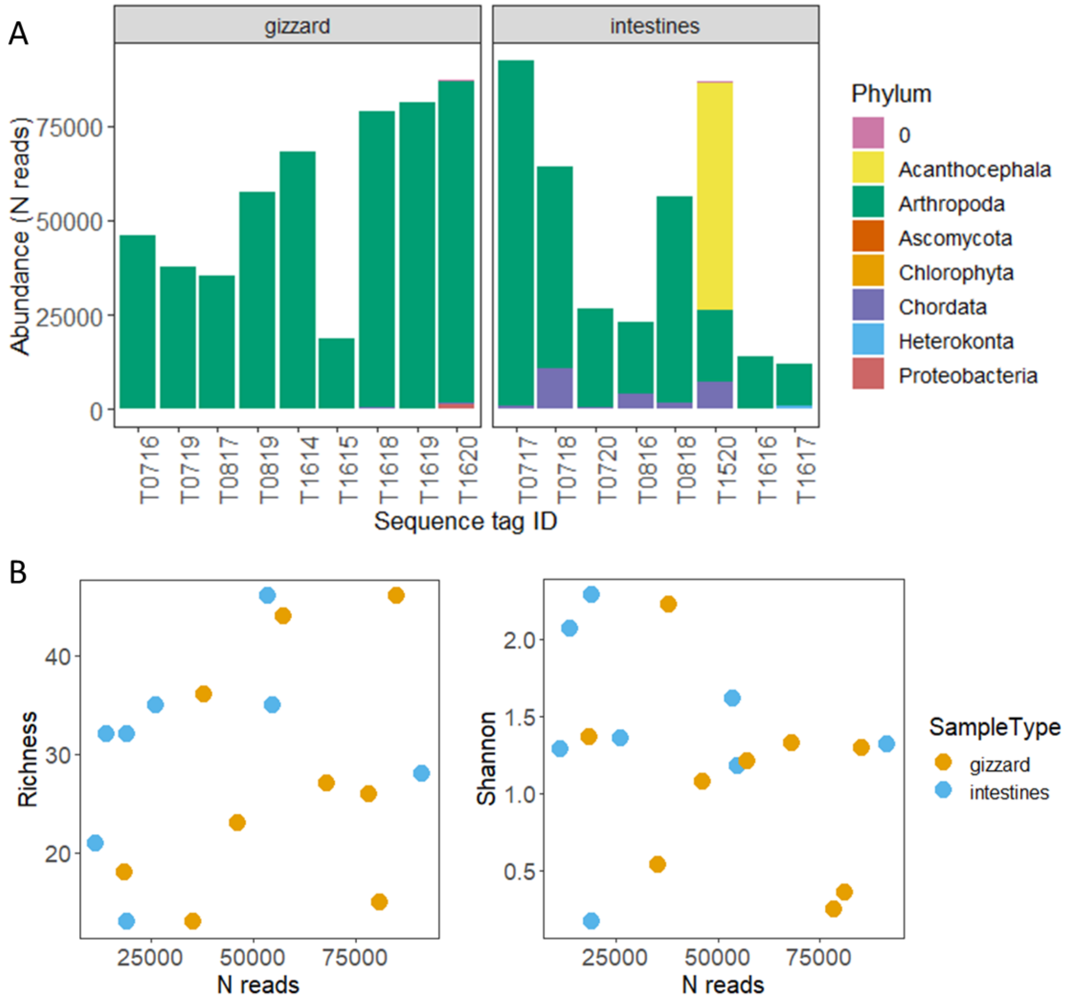


**FIGURE S3-1** Diversity indices of COI barcodes of Test 3. (A) Distribution of reads over all detected phyla for each sequenced sample (incl. a duplicate PCR reaction for one gizzard sample). Zero means unknown phyla. (B) Richness and Shannon diversity versus the total reads per sample. These plots show a reduced data set of only Arthropoda which covered 797,422 reads which is 89.9% of all data (886,749 reads); most data loss (>50,000 reads) was due to parasitic worms found in one intestine sample (phylum: Acanthocephala).

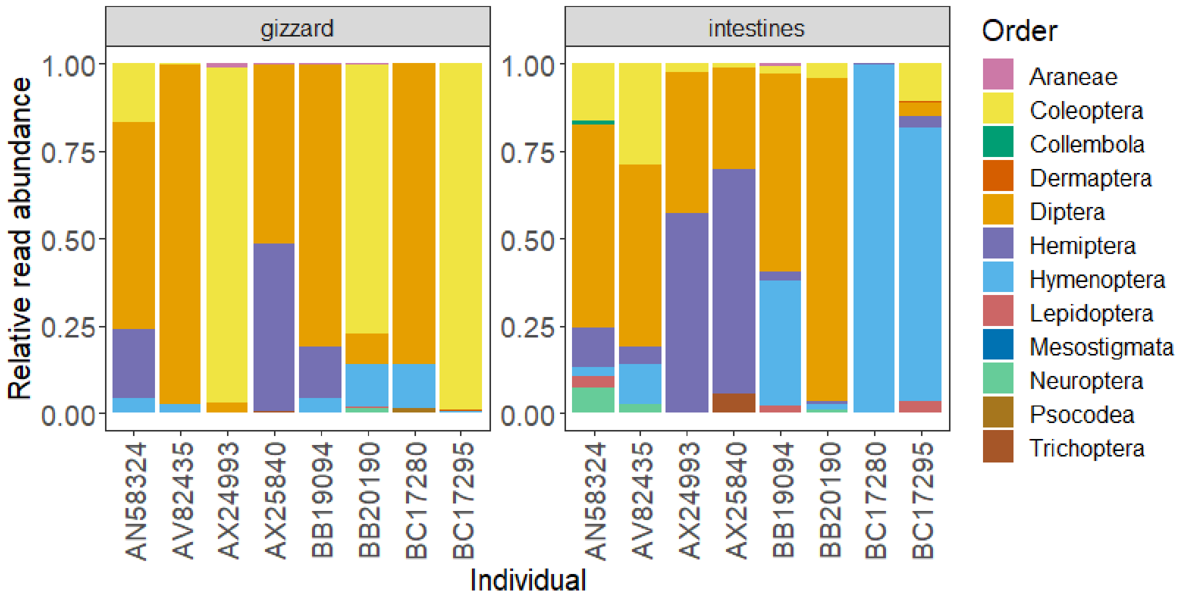


**FIGURE S3-2** RRA of arthropod orders in gizzard versus intestine in each of the eight birds (RingID, i.e. metal band number, indicates each individual). In four of eight individuals, the proportion of reads per order in their gizzard and intestine correlated strongly (R = 0.84-0.98). In the other four birds, either Coleoptera (*N* = 3) or Diptera (*N* = 1) dominated in the gizzard, or Hymenoptera (*N* = 2) or Diptera (*N* = 1) in the intestine (R = -0.08-0.06). Samples were collected from dissected adult male Pied Flycatchers that were found dead in nest boxes.

**TABLE S3-1** Overview of the 20 OTUs detected in the pooled negative PCR controls (NC, sequence ID T0820) of Test 3 and the validation test. Given are GenBank accession numbers of the closest match, the number of reads and the taxonomic assignment

| ID = T0820 | Accession nr. Match GenBank | N reads in NC | Organism | Kingdom | Phylum | Class | Order | Family | Genus |
|---|---|---|---|---|---|---|---|---|---|
| OTU NC_18 | KM449858 | 1 | *Lochmaea capreae* | Animalia | Arthropoda | Insecta | Coleoptera | Chrysomelidae | Lochmaea |
| OTU NC_3 | KU917964 | 7 | *Strophosoma capitatum* | Animalia | Arthropoda | Insecta | Coleoptera | Curculionidae | Strophosoma |
| OTU NC_6 | KX844104 | 5 | *Campylocheta praecox* | Animalia | Arthropoda | Insecta | Diptera | Tachinidae | Campylocheta |
| OTU NC_20 | KM023121 | 1 | *Kleidocerys resedae* | Animalia | Arthropoda | Insecta | Hemiptera | Lygaeidae | Kleidocerys |
| OTU NC_5 | KR897618 | 6 | *Diploplectron sp.* | Animalia | Arthropoda | Insecta | Hymenoptera | Crabronidae | Diploplectron |
| OTU NC_2 | KX665108 | 9 | *Formica sanguinea* | Animalia | Arthropoda | Insecta | Hymenoptera | Formicidae | Formica |
| OTU NC_9 | KR878219 | 3 | *Glypta sp.* | Animalia | Arthropoda | Insecta | Hymenoptera | Ichneumonidae | Glypta |
| OTU NC_12 | MG349058 | 2 | *Pleolophus sp.* | Animalia | Arthropoda | Insecta | Hymenoptera | Ichneumonidae | Pleolophus |
| OTU NC_19 | KX043145 | 1 | *Noctua fimbriata* | Animalia | Arthropoda | Insecta | Lepidoptera | Noctuidae | Noctua |
| OTU NC_17 | LK077046 | 1 | *Apteryx australis* | Animalia | Chordata | Aves | Apterygiformes | Apterygidae | Apteryx |
| OTU NC_15 | GU571396 | 1 | *Ficedula hypoleuca* | Animalia | Chordata | Aves | Passeriformes | Muscicapidae | Ficedula |
| OTU NC_10 | KP719757 | 2 | *Sphyrna lewini* | Animalia | Chordata | Elasmobranchii | Carcharhiniformes | Carcharhinidae | Sphyrna |
| OTU NC_14 | AY049855 | 1 | *Scyliorhinus torazame* | Animalia | Chordata | Elasmobranchii | Carcharhiniformes | Scyliorhinidae | Scyliorhinus |
| OTU NC_7 | AF212180 | 5 | *Triakis semifasciata* | Animalia | Chordata | Elasmobranchii | Carcharhiniformes | Triakidae | Triakis |
| OTU NC_8 | KF808207 | 5 | *Gymnura altavela* | Animalia | Chordata | Elasmobranchii | Myliobatiformes | Gymnuridae | Gymnura |
| OTU NC_11 | KU705517 | 2 | *Rhinoptera bonasus* | Animalia | Chordata | Elasmobranchii | Myliobatiformes | Myliobatidae | Rhinoptera |
| OTU NC_16 | KY176594 | 1 | *Rhinoptera marginata* | Animalia | Chordata | Elasmobranchii | Myliobatiformes | Myliobatidae | Rhinoptera |
| OTU NC_4 | KY176593 | 6 | *Glaucostegus cemiculus* | Animalia | Chordata | Elasmobranchii | Pristiformes/Rhiniformes | Rhinobatidae | Glaucostegus |
| OTU NC_13 | KY176593 | 1 | *Glaucostegus cemiculus* | Animalia | Chordata | Elasmobranchii | Pristiformes/Rhiniformes | Rhinobatidae | Glaucostegus |
| OTU NC_1 | NG_046424 | 10 | *Homo sapiens* | Animalia | Chordata | Mammalia | Primates | Hominidae | Homo |
| **Total** | | **70** | | | | | | | |

**TABLE S3-2** Arthropod taxa community composition in gizzard and intestinal samples of Test 3. Given are quality of match with the GenBank reference sequence (grade) and FOO in either sample type. Only genera occurring FOO ≥ 3 in at least one sample type are shown.

| Order | Family | Genus | Putative species | GRADE | FOO STOMACH | FOO INTEST |
|---|---|---|---|---|---|---|
| Hemiptera | Lygaeidae | Kleidocerys | *Kleidocerys resedae* | 0.982 | 8 | 8 |
| Hymenoptera | Formicidae | Formica | *Formica sanguinea* | 1 | 6 | 7 |
| Diptera | Mycetophilidae | Boletina | *Boletina griphoides* | 1 | 8 | 7 |
| Coleoptera | Curculionidae | Strophosoma | *Strophosoma capitatum* | 1 | 7 | 6 |
| Coleoptera | Chrysomelidae | Lochmaea | *Lochmaea capreae* | 0.902 | 4 | 5 |
| Diptera | Chironomidae | Procladius | *Procladius nigriventris* | 0.998 | 4 | 5 |
| Diptera | Calliphoridae | Pollenia | *Pollenia amentaria* | 1 | 6 | 5 |
| Diptera | Culicidae | Aedes | *Aedes sp.* | 1 | 8 | 5 |
| Diptera | Limoniidae | Limonia | *Limonia nubeculosa* | 0.975 | 2 | 4 |
| Coleoptera | Carabidae | Pterostichus | *Pterostichus oblongopunctatus* | 1 | 4 | 4 |
| Neuroptera | Hemerobiidae | Hemerobius | *Hemerobius micans* | 1 | 1 | 3 |
| Hemiptera | Miridae | Harpocera | *Harpocera thoracica* | 0.998 | 2 | 3 |
| Trichoptera | Limnephilidae | Limnephilus | *Limnephilus auricula* | 1 | 2 | 3 |
| Diptera | Drosophilidae | Phortica | *Phortica sp.* | 0.973 | 3 | 3 |
| Diptera | Tachinidae | Campylocheta | *Campylocheta praecox* | 1 | 4 | 3 |
| Diptera | Chironomidae | Chironomus | *Chironomus sp.* | 1 | 4 | 3 |
| Hymenoptera | Diprionidae | Gilpinia | *Gilpinia virens* | 0.998 | 4 | 3 |

### Supplement 4 of 4

#### Validating the relative contribution of taxa

In the validation test we compared COI metabarcoding data of faecal samples and camera observations. In this supplement additional information is given about the prevailing taxa in faecal samples dominated by a single taxa (**Table S4-1**); the effect of the size-adjustments of prey items (relative to bill size) on the relative abundance of orders in the camera data (**Fig. S4-1)**; the distribution of the number of reads (COI) or prey items (on camera) over the samples and over the unique taxa (**Fig. S4-2**); the observed arthropod diversity and richness on camera (**Fig. S4-3**) and in faecal samples (**Fig. S4-4**); the taxonomic groups per faecal sample on the order level (**Fig. S4-5**); taxonomic assignments and diversity indices for USearch pipeline variants with different filter settings (**Fig. S4-6**); the arthropod communities found in faecal COI data and in the camera data (**Fig. S4-7**) and, lastly, the occurrence of arthropod families in camera recordings and in COI barcodes in (**Fig. S4-8**).

**TABLE S4-1** Domination by a single species and order in COI metabarcoding data: details on the eight of 63 samples that were dominated by a single species (>90% RRA). Shown are RRA of the order dominating the samples, the tentative name of the most common species, the quality of match with the GenBank reference sequence (grade). Note that a taxa may be assigned that has not been described to occur in The Netherlands, which seems the case for *Chrysosyrphus* sp. which is an Artic hoverfly; see https://www.repository.naturalis.nl/document/550168. For the genus *Panzeria* evidence exists that it occurs in The Netherlands, see https://www.repository.naturalis.nl/document/667934.

| YEAR | Sample sequence ID | Order | Proportion orders | Most common species | Grade | N reads common species | Total reads of sample | Proportion species |
|---|---|---|---|---|---|---|---|---|
| 2016 | T0520 | Coleoptera | 0.94 | *Phyllopertha horticola* | 100.0 | 6832 | 7318 | 0.93 |
| 2016 | T1313 | Diptera | 0.99 | *Chrysosyrphus sp.* | 98.6 | 14354 | 14612 | 0.98 |
| 2016 | T0613 | Diptera | 0.95 | *Hybomitra lurida* | 99.1 | 10881 | 11964 | 0.91 |
| 2016 | T0519 | Diptera | 0.96 | *Panzeria rudis* | 100.0 | 4163 | 4466 | 0.93 |
| 2016 | T1315 | Hemiptera | 0.98 | *Stenurella melanura* | 99.8 | 4750 | 5303 | 0.90 |
| 2015 | T0215 | Hymenoptera | 0.97 | *Eridolius sp.* | 96.4 | 15315 | 17573 | 0.87 |
| 2016 | T0515 | Hymenoptera | 0.91 | *Ichneumonidae sp.* | 100.0 | 8351 | 9282 | 0.90 |
| 2015 | T0214 | Lepidoptera | 0.97 | *Colostygia pectinataria* | 100.0 | 1309 | 1563 | 0.84 |

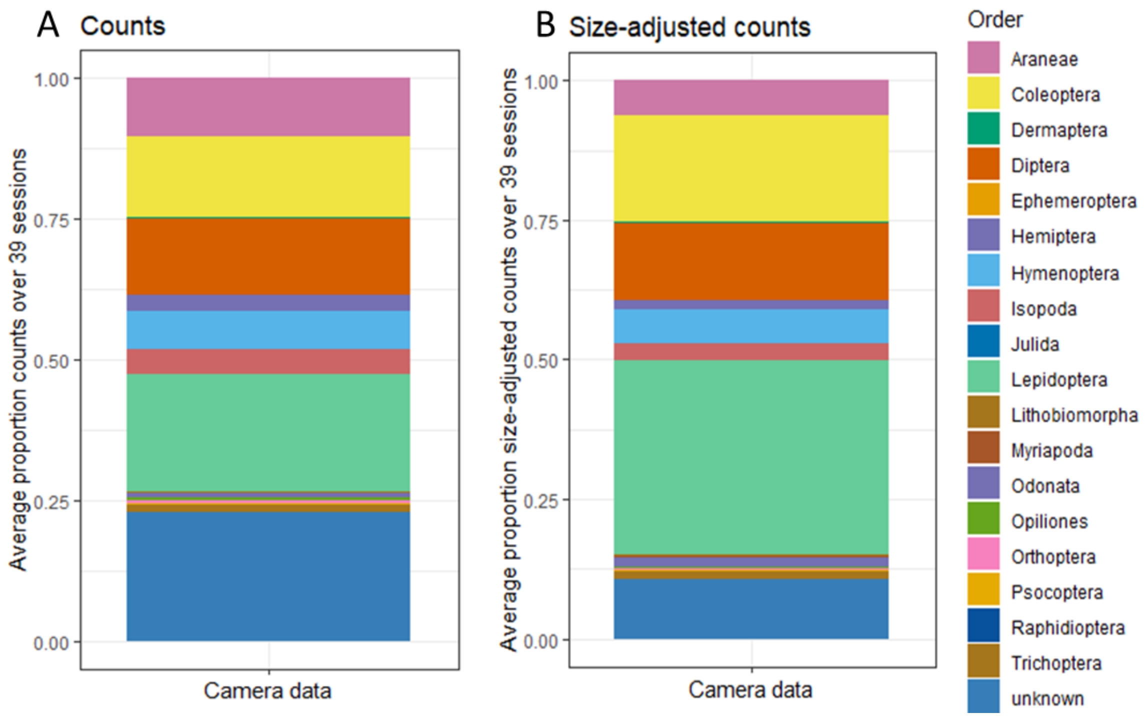


**FIGURE S4-1** The effect of size-adjustments of each prey item (relative to bill size) on the RRA of orders. Compared is the average relative abundance of arthropod orders in the camera records, for (A) raw prey counts; *N* = 7,314, and (B) size-adjusted counts; *N* = 10,458. The average values are based on 39 camera sessions. The stacked bars show the average proportions of each detected order, where the average is the mean proportion over all samples in each year.

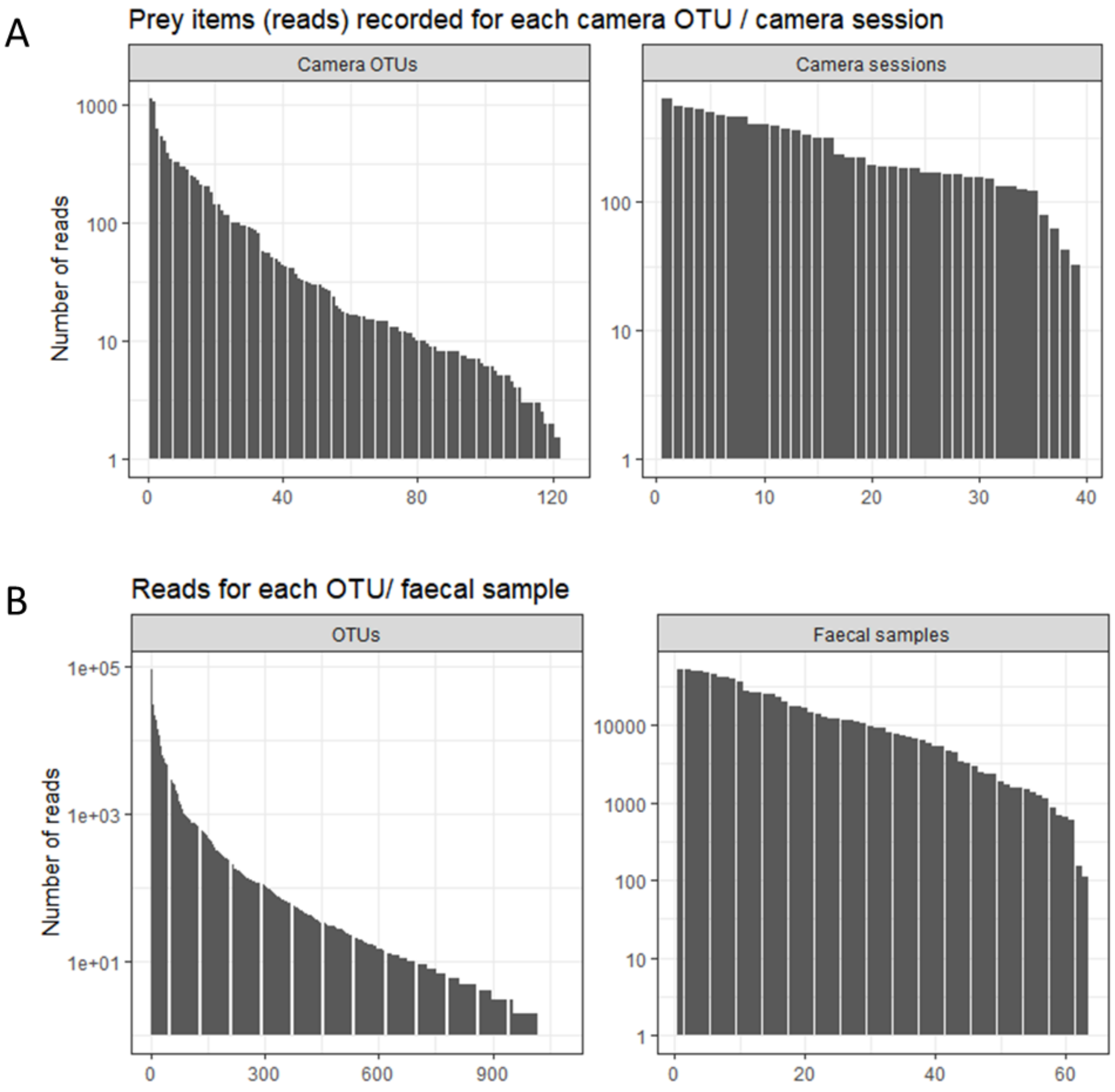


**FIGURE S4-2** Distribution of "reads" in camera recordings (CAM) and in COI barcodes (COI). Given are for CAM data (A): number of size-adjusted counts ("reads") for each of the 124 unique prey items (camera OTUs) (left panel) and the number of size-adjusted prey counts ("reads") recorded in each of the 39 camera sessions (right panel). For the COI data (B), we plotted number of reads of 1,083 OTUs (left panel) and 63 samples (right panel).

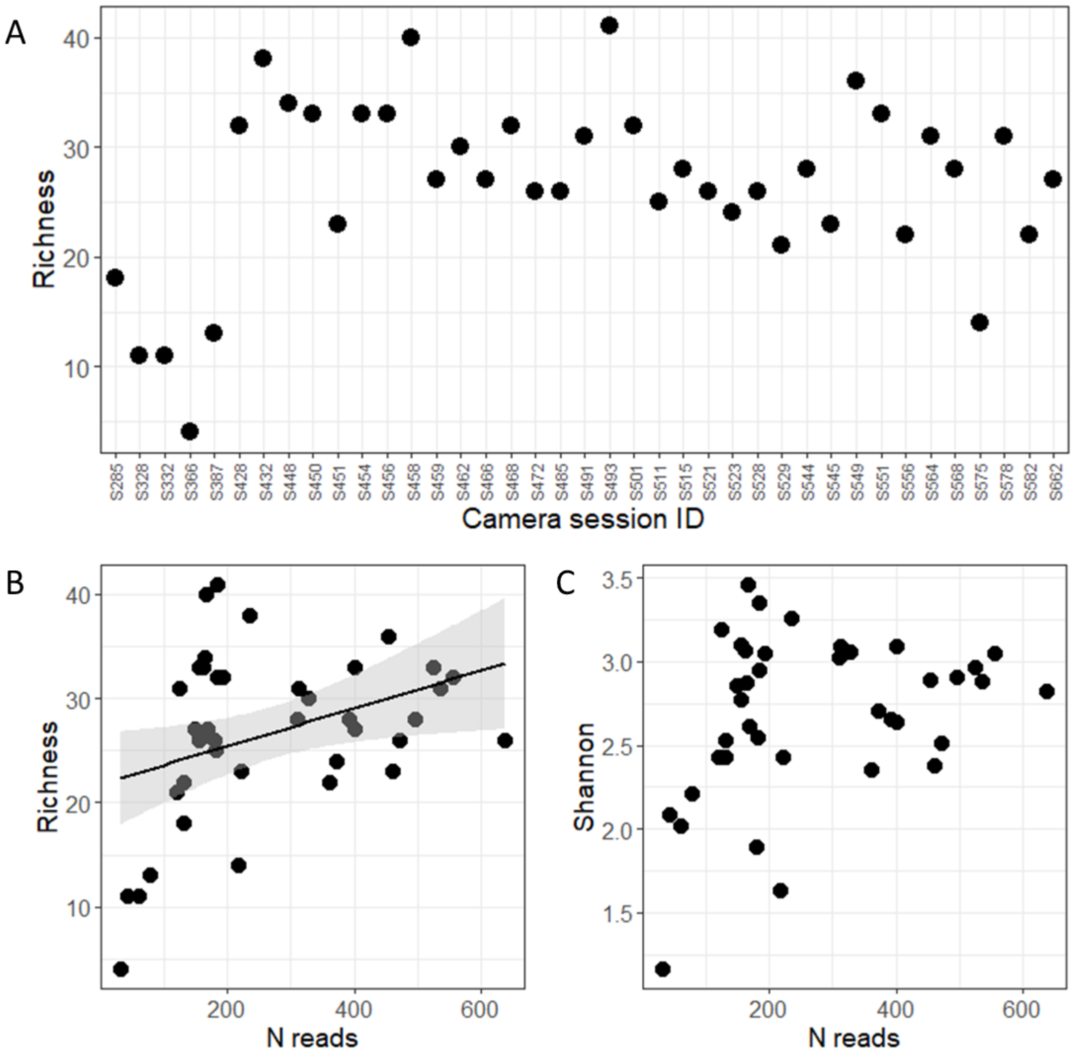


**FIGURE S4-3** Diversity indices of the camera observations of the validation test. The camera data contained four arthropod classes: Arachnida, Insecta, Diplopoda and Malacostraca which are all included in here. (A) Richness per sample. (B) Richness - number of camera OTUs detected per sample and the total number of size-adjusted counts (“reads”) per sample. (C) Shannon diversity and the total number of size-adjusted counts (“reads”) per sample.

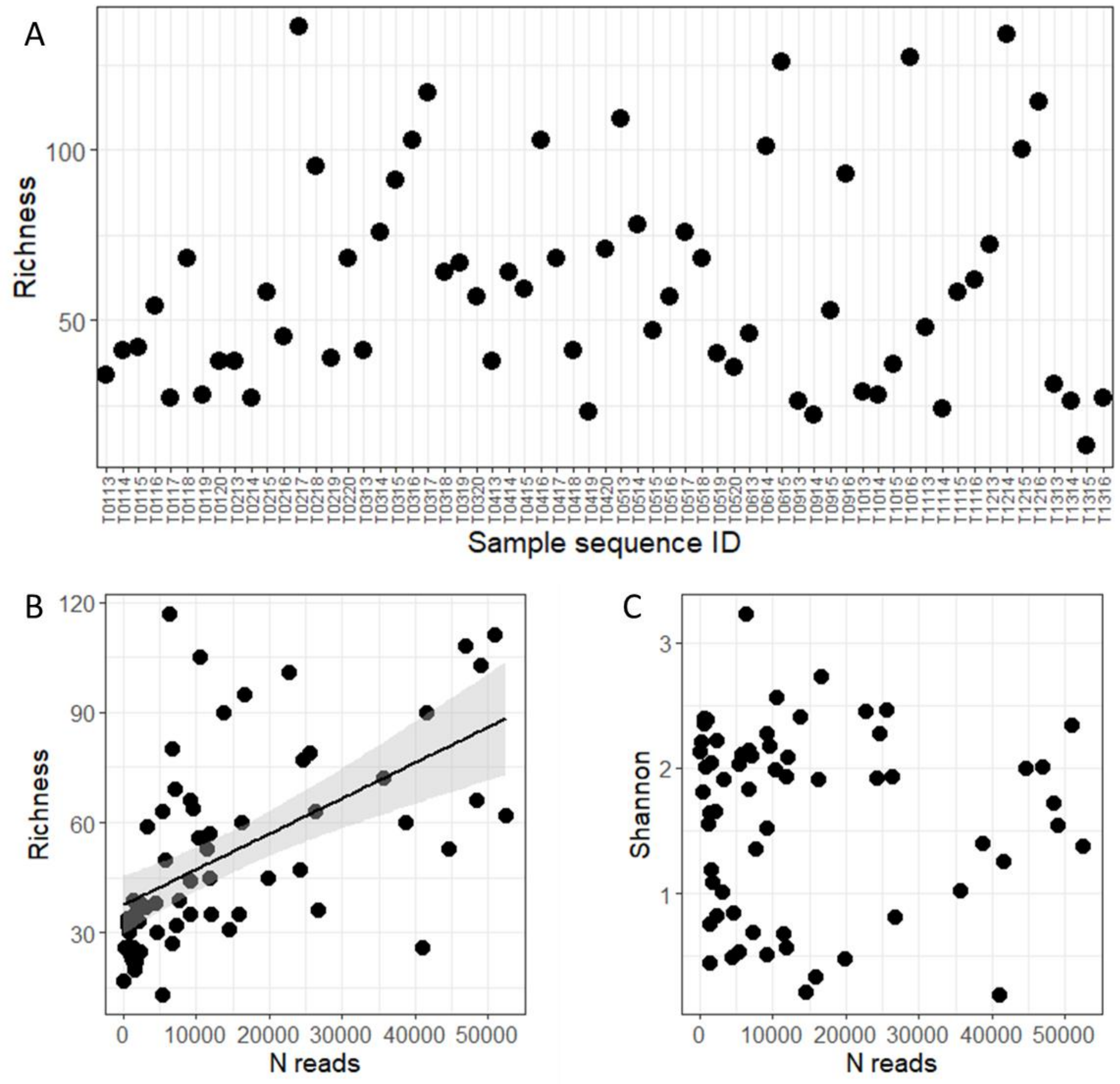


**FIGURE S4-4** Diversity indices of the COI barcodes of validation test. (A) Number of OTUs detected per sample for all data (911,947 paired reads) which included 43 classes of four Kingdoms (Animalia, Bacteria, Plantae and Protista) and unclassified taxa (eukaryote or protozoa). (B) Number of arthropod OTUs detected per sample plotted against total number of paired reads per sample. (C) Shannon diversity plotted against total number of paired reads per sample. For plot B and C, the reduced data set was used (897,315 paired reads) which included five arthropod classes (Arachnida, Insecta, Chilopoda, Diplopoda and Malacostraca) which covered 98.4% of all data.

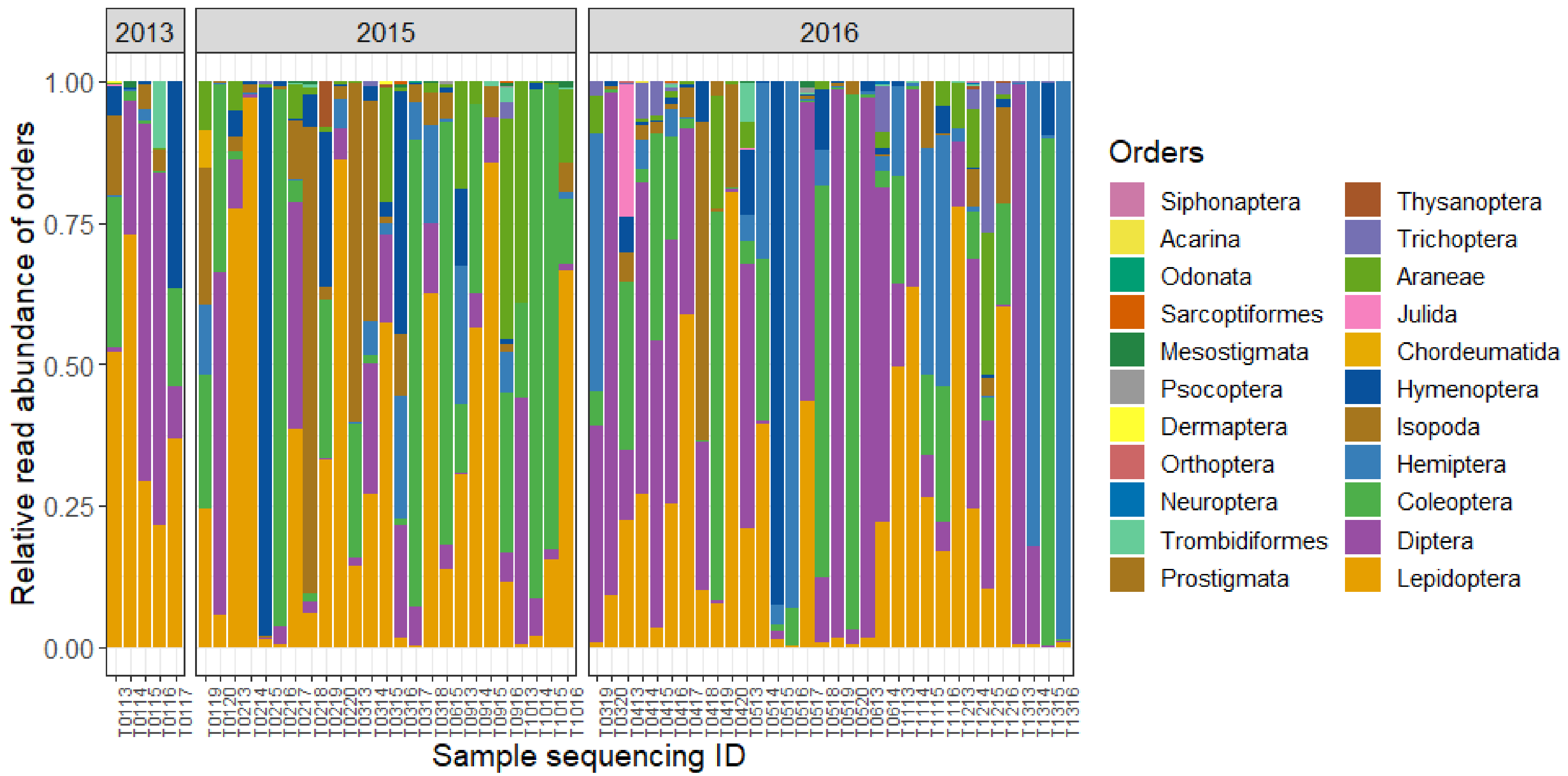


**FIGURE S4-5** Relative read abundance (RRA) of arthropod orders in each sample, to illustrate potential domination by a single order (and species) in COI metabarcoding data. The stacked bars show the proportions of each detected order. Information on detailed taxonomy of samples dominated by a single order is given in **TABLE** S4-1.

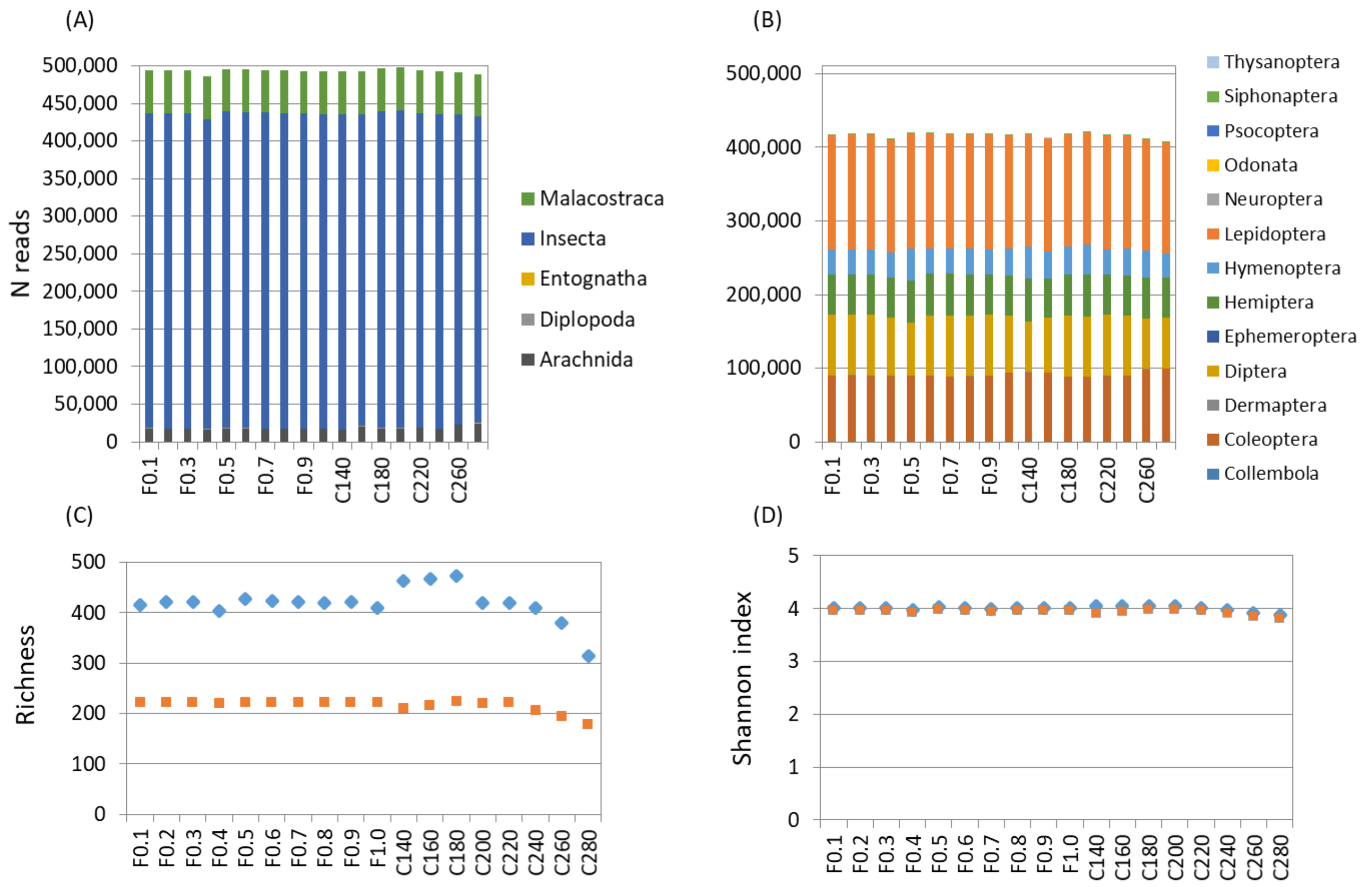


**FIGURE S4-6** Taxonomic assignments and diversity indices for USearch pipeline variants with different filter settings (F0.1 to F1.0; 10 categories) and trimming cut-offs (C140 to C280; 8 categories where number refers to bp). Given are (A) absolute number of reads assigned to arthropod classes ($N$ = 5) and (B) to Insecta orders ($N$ = 13); rare taxa are listed in legend but not visible in plot. Family levels are not shown (see text). Diversity is shown as (C) richness and by (D) the Shannon index, for all data (blue symbols) and for the 99% dataset (orange symbols) which excluded species occurring <0.01%.

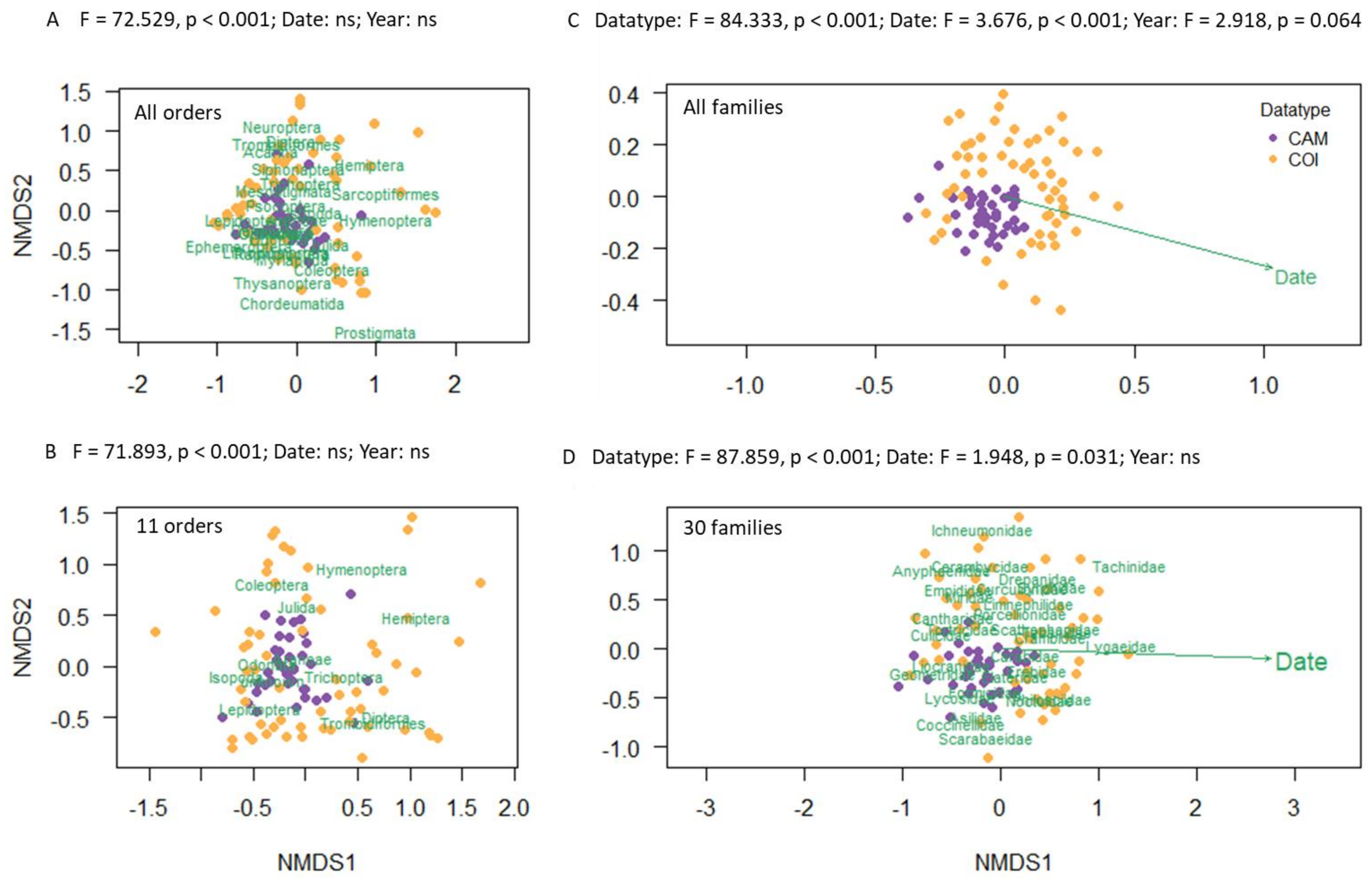


**FIGURE S4-7** Non-metric Bray-Curtis ordination plots comparing communities in camera observations (CAM) and COI metabarcodes (COI). Each sample is represented by a data point. The placement of taxa is indicated in green. A: all arthropod orders detected in both datasets, excluding the category arthropod order “unknown”. B: 11 most common orders plus the category order “unknown”. C: all 153 detected arthropod families in both datasets (145 in COI and 8 more on camera). D*:* top 30 most common arthropod families. The arrow indicates that community varied significantly with date of the season.

*Guide to the interpretation of Fig. S4-8:*

In **Diptera**, the most common family, Tabanidae (horse flies), and the most common species (*Hybomitra lurida*, broad-headed horsefly) was the same in camera and COI data, and also the relative abundances of Syrphidae (hoverflies), Scathophagidae (dung flies), Culicidae (mosquitos) and Asilidae (stiletto flies) were similar. Empididae (dance flies), Tachinidae (true flies) and Ephydridae (shore flies) were commonly detected in COI but not or rarely on camera. Rhagionidae (snipe flies) were often on camera but very rare in COI (< 1% of reads). In **Lepidoptera**, the most common taxa in both datasets were Noctuidae (owlet moths), Geometridae (geometer moths), Depranidae (hooktip moths), Tortricidae (leafroller moths) and Erebidae (erebids moths). Not recorded on camera were Ypsolophidae (a.o. feather-horn moths) and Crambidae (grass or granite moths). In **Coleoptera**, the overlapping common families were Elateridae (click beetles), Curculionidae (weevils), Cantharidae (soldier beetles) and Scarabaeidae (garden chafers, present with several sequence variants). Carabidae (ground beetles) and Cerambycidae (longhorn beetles) were common in COI, but rare on camera, while Tenebrionidae (darkling beetles) and Coccinellidae (ladybirds) were more common on camera than in COI (< 1% of reads). In **Hymenoptera**, **Hemiptera**, and **Araneae** the common families detected on camera and in COI did not strongly overlap much; in these taxa identification from the pictures was mostly impossible by our observers, and therefore in the camera records >10% of prey observations were logged as arthropod family “unknown” (see Fig. 4 in the main text).

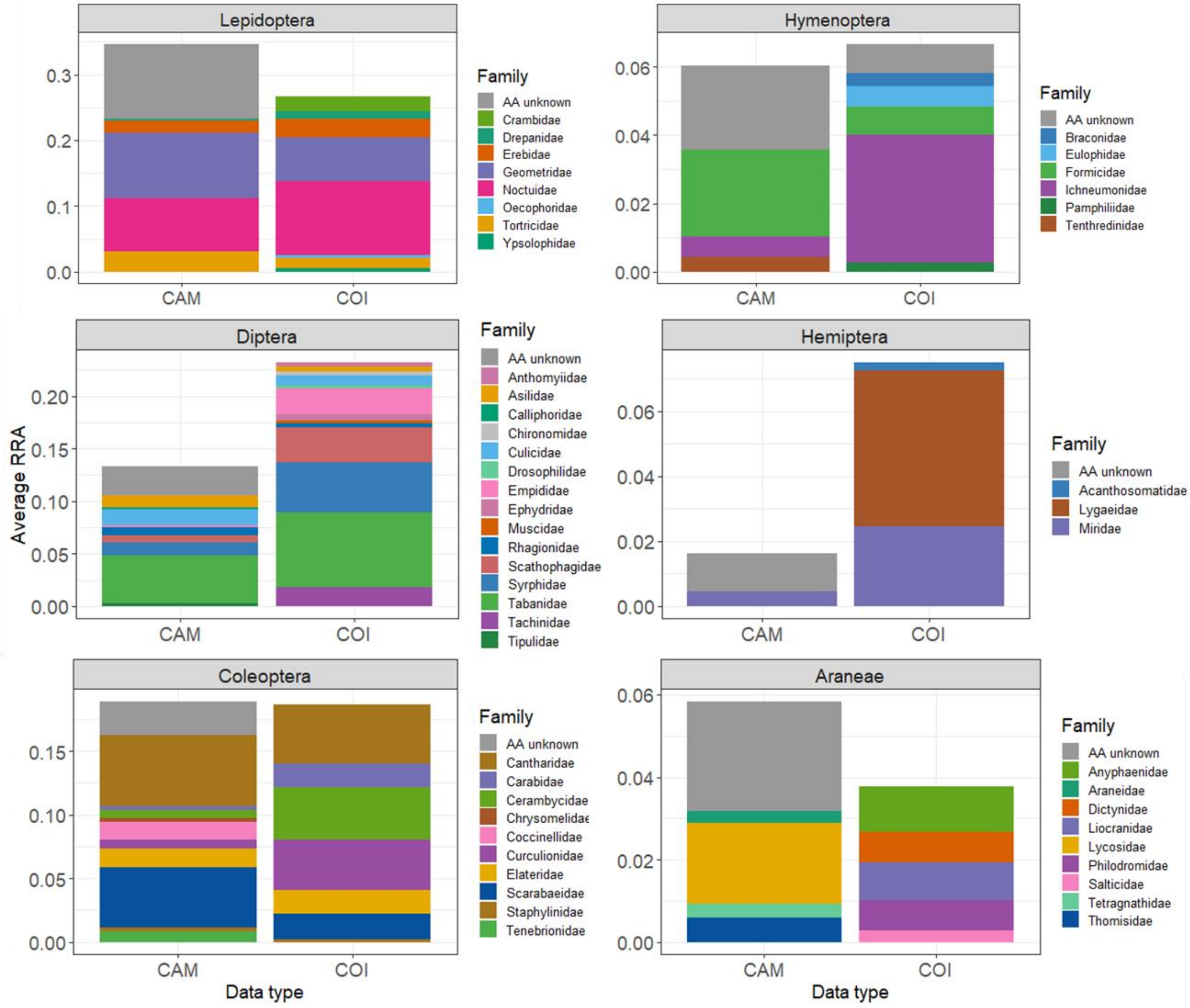


**FIGURE S4-8** Comparison of the occurrence of arthropod families in camera recordings (CAM) and in COI barcodes (COI). The dataset was reduced to the six most abundant orders which were the same in both datasets. Shown are the average relative read abundances (RRA) of common arthropod families per order. “Common” was defined as average read abundance >0.2% for combined data. Because the dataset was pruned the proportions do not add up to 1.

References

Brown, D. S., Jarman, S. N., & Symondson, W. O. C. (2012). Pyrosequencing of prey DNA in reptile faeces: Analysis of earthworm consumption by slow worms. *Molecular Ecology Resources*, *12*(2), 259–266. doi:10.1111/j.1755-0998.2011.03098.x

Folmer, O., Black, M., Hoeh, W., Lutz, R., & Vrijenhoek, R. (1994). DNA primers for amplification of mitochondrial cytochrome c oxidase subunit I from diverse metazoan invertebrates. *Molecular Marine Biology and Biotechnology*, *3*(5), 294–299.
